# A high-density lineage tree reveals dynamics of expression differences accumulation in nondifferentiating clonal expansion

**DOI:** 10.1101/2021.11.24.469964

**Authors:** Feng Chen, Zizhang Li, Xiaoyu Zhang, Peng Wu, Wenjing Yang, Xiaoshu Chen, Jian-Rong Yang

**Affiliations:** Department of Biomedical Informatics, Zhongshan School of Medicine, Sun Yat-sen University, Guangzhou 510080, China; Department of Medical Genetics, Zhongshan School of Medicine, Sun Yat-sen University, Guangzhou 510080, China

**Keywords:** Cell lineage tree, Expression noise, Homeostasis

## Abstract

Differences in gene expression levels among genetically identical cells naturally accumulate during cellular proliferation, forming the basis of expression noise or differentiation. Nevertheless, how transcriptome-wide noise accumulation is constrained to maintain homeostasis during continuous cell divisions has remained largely unresolved. We developed a novel method named “single-cell transcriptome and dense tree” (STADT) to simultaneously determines the transcriptomes and lineage tree of >50% single cells in a single-cell-seeded colony. This lineage tree revealed gradual accumulation of transcriptome differences that became saturated upon four cell divisions, reduced expression noise for sub-tree/sub-colonies closer to inferred expression boundaries, and transcriptionally modulated co-fluctuations among genes. These results collectively showed, for the first time, constrained dynamics of expression noise in the context of cell division.

## Introduction

Genetically identical cells often display significant differences in the expression of individual genes ^1, 2^. Such expression differences are generally considered to be among the most fundamental features underlying the vast diversity and complexity of life ^2, 3^. There are two tightly-connected manifestations of expression differences: expression noise and cellular differentiation. Here, expression noise refers to expression differences among cells growing in the same environment, but has yet to create significant functional/morphological alteration for the cells. Gene expression noise is subject to natural selection ^4, 5^ and tight genetic regulation ^6^. In addition, expression noise is considered a trigger for differentiation ^7, 8^, which involves changes in (population-average) expression levels stabilized by epigenetic regulation and is the basic biological process leading to the development of different functional types of cells (i.e., cell identity).

Expression noise is clearly influenced by cell divisions. However, our current understandings for the regulation of expression noise at the transcriptome level is usually derived from static snapshots of a population of cells and is about the molecular regulation of transcription or transcript degradation ^6, 9–11^, but has been less studied in the context of cell divisions ^12, 13^. Specifically, does expression noise propagates as a single cell continuously divides to generate a population of cells (i.e., single-cell clones) while maintaining the homeostasis? In theory, stochastic partition of molecules could on the one hand exaggerate expression noise and therefore increasingly threat homeostasis maintenance ^13^. On the other hand, timely division could also constrain expression noise, as the expression difference between sister cells should in theory gradually increase after their division from the mother cell ^14^. To resolve the role of cell division in regulating expression noise, one should ideally trace all the cell divisions occuring in the expansion of the clonal colony, while measuring the transcriptomes of all the cells. Unfortunately, this is currently impossible as transcriptome measurements routinely involve RNA extraction and therefore killing the cells.

One potential strategy for overcoming this problem is the simultaneous determination of single-cell transcriptomes and lineage trees for single-cell clones. Here a lineage tree is a record of (ideally) all cell lineage division events along the growth of the single-cell clone up to the time-point of transcriptome measurement, which therefore constitutes a hierarchical tree-like structure among single cells as defined by their kinship. A sub-tree of the lineage tree is similarly defined, but just for the growth of a sub-colony that ends either at or before the time-point of transcriptome measurement (**Figure S1**). The advantage of such data is the large number of samples spanning many stages (corresponding to the number of cell divisions) of clonal expansion under the assumption that any sub-tree rooted at intermediate cells of the tree represents an unbiased sample of clonal expansion (**Figure S1**). This assumption was supported by the observation that the correlation of expression levels among individual cells within the same lineage tree depends only on the genealogical distance along the tree and independent of the position relative to the root ^15^. The recent combination of single-cell RNA-seq (scRNA-seq) with genome editing (e.g., single-cell (sc) GESTALT ^16^ and CARLIN ^17^) has provided a unique opportunity towards this goal. However, scGESTALT-like techniques usually give rise to sparsely sampled (in terms of the number of cells within the colony) lineage trees due to various technical limitations, such as a high rate of long-deletions in the lineage barcode and a low recovery rate of lineage barcodes via scRNA-seq, thereby preventing their application in resolving the propagation of expression noise during clonal expansion.

In the current study, we measured the single-cell transcriptomes and generated lineage trees of single-cell clones consisting of up to ∼5000 HEK-293 cells using a newly developed experimental pipeline referred to as the “*s*ingle-cell *t*ranscriptome *a*nd *d*ense *t*ree” (STADT) method. Utilizing inducible Cas9 and constitutively expressed lineage barcode, with optimized editing efficiency in terms of highly sequential editing events with a relatively low frequency (∼8%) of long-deletions, STADT determined the lineal positions of up to 93% of cells captured by 10x Chromium and >50% of the cells in the original colony. More importantly, up to 92% of the tips (i.e., leaves) in the reconstructed lineage trees are single-cell tips, rather than clusters of cells carrying identically edited lineage barcodes. This lineage tree facilitated analyses of expression noise in the context of cell division with an unprecedented resolution and revealed noise saturation after four divisions, decelerated noise accumulation in sub-trees rooted with cells showing expression levels near an upper limit, and co-fluctuation among gene pairs whose dosage balance is functionally essential. Our results therefore not only represent an novel effort in the genetic reconstruction of lineage trees towards higher coverage of single cells, but also reveal the cellular constraints on the accumulation of expression noise with respect to cell division/clonal expansion, a previously underappreciated mechanism of homeostatic maintenance in the face of stochastic expression noise.

## Results

### STADT reconstructs a dense lineage tree for a single-cell clone

Our goal was to construct a dense lineage tree of a single-cell clone grown *in vitro*, in which the transcriptomes at the tips were measured, as we reasoned that the high sampling density of cells should help in the approximation of divergence times between cell pairs in the unit of divisions. To this end, the simultaneous measurement of the single-cell transcriptomes and the lineal relationships of as many cells as possible are required. Recently developed scGESTALT technology ^16^ has shown potential for generating such data via the combination of the cumulative editing of genome-integrated lineage barcode and scRNA-seq, albeit with a clear pitfall in terms of the density of sampled cells within the population/lineage tree (see below). We therefore adopted the scGESTALT protocol with four major revisions in STADT. First, we designed four single guide RNAs (sgRNAs) and a 432 base pair-long lineage barcode (hereafter referred to as the “STADT barcode”) containing 13 target sites with up to 3 nucleotide mismatches with the sgRNAs (**Figure 1A** and **Table S1**, details in **Methods**). Mismatches were introduced to reduce the editing efficiency ^18^ and thus the probability of multi-site long deletions. Second, in addition to the STADT barcode and sgRNAs, Cas9 driven by an inducible CuO promoter (CuO-Cas9) and the corresponding suppressor CymR was inserted into the HEK-293 genome. Cas9 could therefore be expressed only in the presence of cumate, allowing the synchronized initiation of clonal expansion and lineage barcode editing (**Figure 1A**). Third, we monitored the expression levels of transgenes (**Figure S2** and **Table S2**), the percentage of living cells and the overall morphology of the single-cell colonies (**Figure S3**, see **Methods**), and the colonies that were most likely to exhibit a high proportion of surviving cells with edited STADT barcodes (hereafter “STADT alleles”) were chosen for the subsequent digestion (see **Methods**). We validated the STADT alleles for sufficient diversity and desirable resolution for lineage tree reconstruction by Sanger (**Figure S4**) and high-throughput sequencing (**Figure S5**. See also **Methods**). Fourth, we developed a four-step digestion protocol to minimize the number of cells killed during digestion such that the fraction of surviving cells subjected to 10x Chromium library preparation was maximized (**Figure S3C**, see **Methods**).

**Figure 1.**
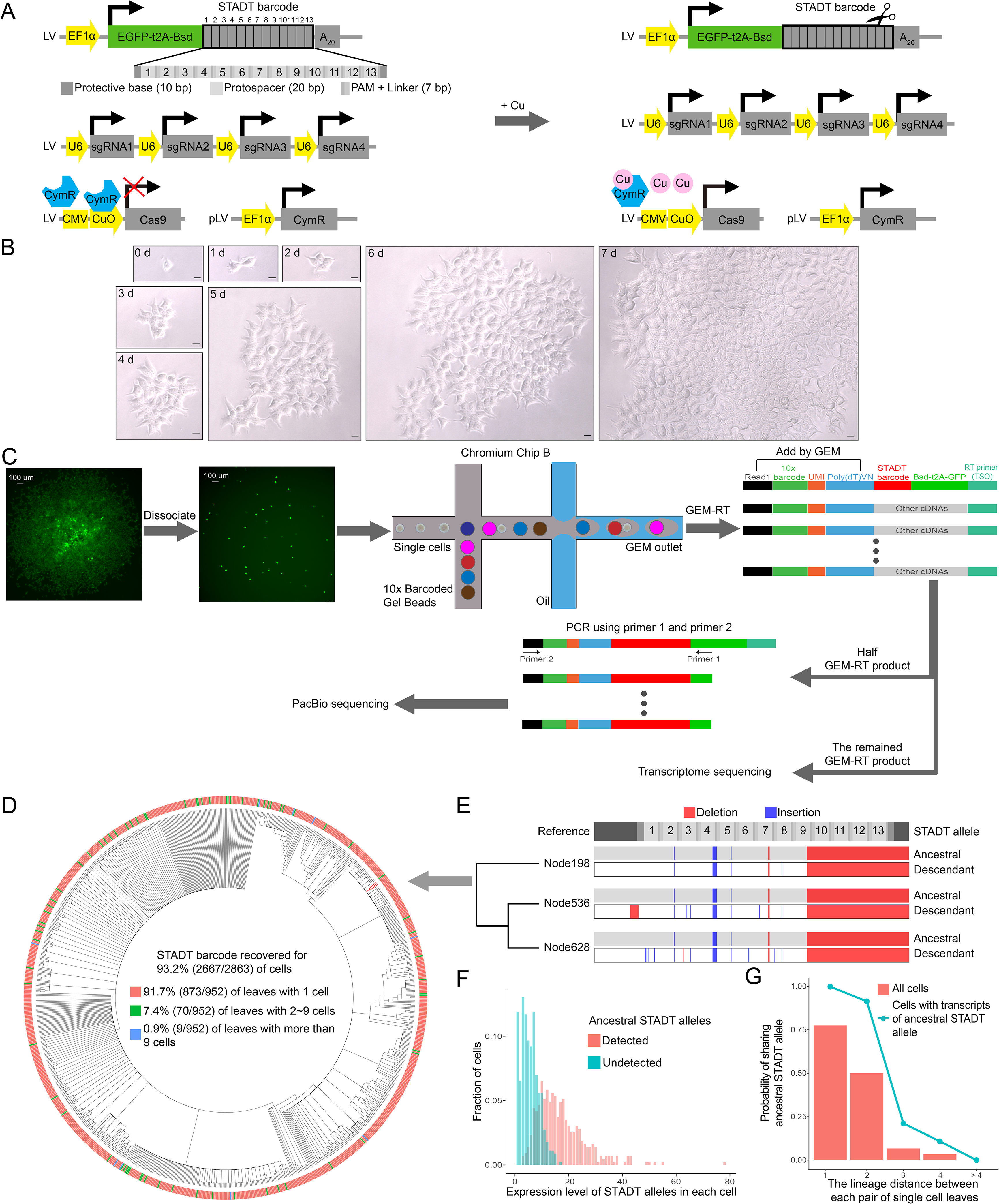
The STADT method reliably determined the lineage tree for majority of the cells within a single-cell clone. **(A)** STADT components designed for lineage tracing in HEK-293 cells. The STADT barcode and sgRNAs were randomly inserted with a lentivirus vector system into the genome of an HEK-293 cell line with inducible Cas9 expression, which already contained Cas9 and CymR expression cassettes (see Methods, and Figure S2). The addition of cumate (right) to the culture medium resulted in rapid sequestration of the CymR protein, preventing its binding to the operator (CuO) and thereby enabling the expression of Cas9. **(B)** The morphology of a single-cell colony during seven days of clonal expansion. The scale bar indicates 20 µm. **(C)** Schematic diagram showing the preparation and the single-cell sequencing of a colony with 10x Genomics Chromium technology (see Methods). **(D)** The reconstructed lineage tree of a single-cell colony. A total of 2863 cells were captured by scRNA-seq, among which 2667 had STADT barcodes recovered by PacBio-CCS. The STADT barcodes were used to construct a maximum likelihood tree (see Methods). Some cells had identical STADT barcodes and were collectively represented as a single leaf in the tree. The number of cells represented by each leaf of the tree is shown by the color tile at the outer ring of the circularized tree, using the colors indicated in the legend at the center of the tree. **(E)** An example of cells with a common ancestral STADT allele. This particular subtree is indicated by the red-colored branches in the full tree shown in panel d. The structure of the STADT barcode is indicated on top by gray gradients similar to that in Figure 1a. Insertion and deletions are respectively indicated by blue and red strips. **(F)** Cells with high expression of the STADT barcode tended to contain the ancestral STADT allele. **(G)** For a pair of single-cell leaves, with increasing lineage distance in the tree (*x* axis), the probability of finding a common ancestral STADT allele between the two cells decreased. This pattern was consistent within all cells (red bars) or within cells containing transcripts of the ancestral STADT allele (cyan line). The lineage distance between a pair of cells was defined as the larger (between the two cells) number of detected divisions since the most recent common ancestor.

Applying the STADT method, we digested three single-cell clones of approximately 5,000 ∼ 10,000 cells each (samples “A”, “B” and “C”, **Figure S6**) and loaded them individually into a GemCode Single Cell Instrument (10x Genomics) to generate cDNAs corresponding to the transcripts of each single cell. Each resulting library was split into two halves: one half was subjected to NovaSeq-based RNA-seq for single-cell transcriptomes, and the other half was subjected to amplification of the STADT barcode, followed by PacBio Sequel-based HiFi sequencing of the STADT barcode (**Figure 1C**, see also **Tables S3** and **S4**). A lineage tree was then constructed for each sample based on the STADT alleles via the maximum likelihood method (**Methods**) for the cells with both single-cell transcriptome and STADT barcode determined. We found that the lineage tree of sample A, containing 2667 single cells (**Figure 1D**), showed a low frequency of long deletions (8.8% of all unique STADT alleles), a high fraction of captured cells (93.2% of cells captured by RNA-seq, >50% of cells in the colony), a high fraction of single-cell tips (91.7% of all tips) and a deep lineage tree (tree depth was 18 divisions) (**Tables S4** and **S5**), and we therefore decided to focus on this sample in the downstream analyses.

To verify that the overall hierarchical relationships among the cells captured in the lineage tree were largely correct, we extracted the ancestral STADT alleles retained by descendent cells in the form of yet-to-decay transcripts (an example provided in **Figure 1E**), which were particularly abundant in cells with high expression of STADT barcodes (**Figure 1F**. See **Methods**). The lineage relationship of a pair of sister cells is more likely correct if shared ancestral STADT alleles can be found in both cells. In addition, due to RNA decay, the probability of observing shared ancestral STADT alleles should decrease (relative to that in sister cells) in first-cousin cells and decrease further in second-cousin cells. This situation was indeed observed in the reconstructed lineage tree (**Figure 1G**). In combination with the aforementioned results suggesting a fine resolution of early division events (**Figure S5K**), we concluded that a lineage tree with a high density of cells within a single-cell clone was obtained for sample A, with reasonably well-resolved hierarchical relationships observed among the cells. Nevertheless, we wish to emphasize that although STADT outperformed scGESTALT-like techniques in terms of resolution (of the hierarchical relationships among the cells) and density (of cells within the single-cell clone), the STADT-based lineage tree is still too coarse grained for detailed developmental analyses, such as those conducted in *Caenorhabditis elegans* ^19–21^.

### Signature of noise accumulation and saturation during clonal expansion

We then examined noise accumulation during clonal expansion in the reconstructed lineage tree. We chose to use the depth of the sub-tree, defined as the length (in units of the number of cell divisions) of the longest branch in the sub-tree (**Figure 2A**), as a quantitative metric for the stage of clonal expansion a sub-tree is in (see **Methods**). The divergence time between any pair of cells was then defined as the depth of the sub-tree rooted at the most recent common ancestor. Under the assumption that the sub-trees were all unbiased samples of clonal expansion (**Figure S1**), we can compare the level of expression noise in different stages of clonal expansion. As a demonstration of this approach, we randomly chose three sub-trees with depths of 1, 2 and 3 (**Figure 2B**). The expression noise among the single-cell leaves of these subtrees as measured by coefficient of variation (CV) was calculated using the 100 genes with the highest expression (according to the average across the full tree) (**Figure 2C**), which revealed a progressive increase in expression noise with clonal expansion. Indeed, deeper sub-trees showed greater variation according to a principal component analysis using all genes (**Figure 2D**). To globally gauge the accumulation of expression noise during clonal expansion, we compared the divergence time and the normalized expression difference (Euclidean distance of the expression of all genes) for all pairs of single-cell leaves (**Figure 2E**). We again found evidence of greater expression differences in cell pairs with longer divergence times (Spearman’s rank correlation coefficient _ρ_ = 0.31, *P* < 10^-8^).

**Figure 2.**
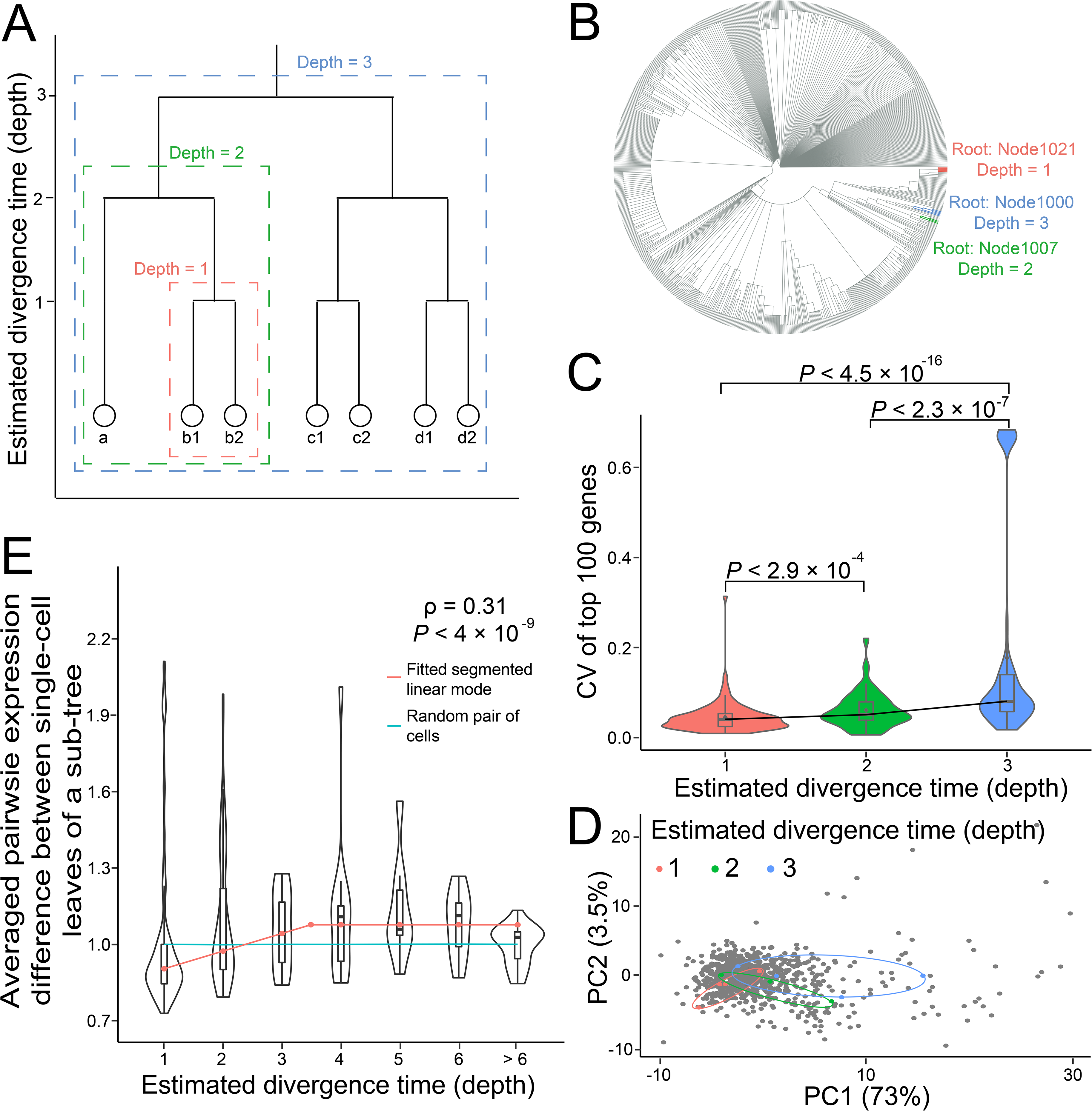
Signatures of noise accumulation and saturation. **(A)** Schematic diagram for the estimation of divergence times between pairs of cells, which was approximated by the depth of the sub-tree rooted by the most recent common ancestor of the focal pair of cells. For example, the divergence time between cell “b1” and cell “b2” is 1, as the sub-tree rooted by their most recent common ancestor has a depth of 1 (red box). Similarly, the divergence time between “a” and either “b1” or “b2” is 2 (green box). The divergence time between any of “a”, “b1” or “b2” and any of “c1”, “c2”, “d1” or “d2” is 3 (blue box). **(B)** Three sample sub-trees with depths of 1 (red), 2 (green) or 3 (blue) were chosen from the full lineage of the single-cell colony and further analyzed as shown in panels c and d. **(C)** Then, the CV of the expression levels of the top 100 most highly expressed genes was calculated among the single-cell leaves of the three sample sub-trees, and the distribution of the 100 CVs is shown in violin plots, revealing a gradual increase in expression noise. *P* values of Wilcoxon signed-rank tests between sub-trees are indicated on top. **(D)** The accumulation of expression noise is also supported by the greater span (ovals) observed among the leaves (dots) of deeper sub-trees (colored as in panel b) when the transcriptional profiles (of all genes) of all single-cell leaves were visualized by principal component analysis (PCA). The percent variance explained is indicated for principal components 1 (PC1, *x* axis) and 2 (PC2, *y* axis). **(E)** All pairwise expression differences (Euclidean distance of the expression of all genes) were calculated for the single-cell leaves and then normalized via division by the average pairwise distance of 1,000 random pairs of cells, such that the expected expression difference between any pair of cells was equal to 1 (the cyan line). The distribution of expression differences between pairs of cells with the same estimated divergence time is shown as a violin plot. Comparison among cell pairs of different divergence time revealed the accumulation and saturation of expression differences or noise. Spearman’s rank correlation coefficient for the original normalized expression differences is indicated in the top right corner along with the corresponding *P* value. The red line indicates the segmented linear model fitted with the normalized expression difference.

The gradual accumulation of expression noise should eventually break the homeostasis of the population, which is in apparent contrast to common observations for HEK-293 cell line and suggests that noise accumulation should saturate at some point. To verify this notion, we fit the growth of expression noise with divergence time by a segmented linear model assuming eventual noise saturation ^22^ (hereafter “saturation” model. **Figure 2E**, red line). Indeed, we found that this saturation model significantly outperformed other linear models assuming either unlimited noise accumulation along clonal expansion (“unlimited” model) or complete independence between noise and divergence time (“independent” model) (**Table S7**). According to the saturation model, 83% of the expression differences among cells are attributable to expression fluctuations within a cell cycle, which is largely consistent with previous findings ^23, 24^. The estimated saturation time was at the divergence time of 3.49 suggesting that four cell divisions were required (either temporally or functionally) to realize the full range of gene expression noise. This observation further indicates that cellular inheritance of the transcriptome on average lasts for at least three generations/divisions.

To further resolve this pattern of noise saturation, we analyzed the noise accumulation of individual genes. Consistent with the transcriptomic result, we found that out of the 710 genes with sufficient data (**Table S7**) to fit all three models (saturation, independent and unlimited), 492 were best described by the saturation model. We found significant enrichment of genes functioning in mitochondria and translation/ribosome (**Figure S7**) among these 492 genes with apparent signature of noise saturation. This is not unexpected because proteins of (nuclear-encoded) mitochondria and ribosome-related genes are on the one hand major factors driving extrinsic noise^25–28^, and on the other hand heavily influenced by stochastic partition at cell division, as they tend to segregate^29^ (in the form of mitochondrion and polyribosome, respectively). The combination of these two features suggests that noise constraints for mitochondria and ribosome-related genes should have been favored by natural selection to reduce the global expression noise^30^. Furthermore, there is significant among-gene variation in the saturation time. For example, some genes, such as the Ribosomal Protein L11 (*RPL11*) gene, showed noise saturation at the divergence point of only two divisions. Other genes, such as glutathione peroxidase 4 (*GPX4*), must undergo 5 divisions to achieve the full range of expression noise. Interestingly, the variation in saturation time seemed unexplainable by previously reported^31^ gene-specific mRNA turnover rates in HEK-293 cells (**Figure S8**). These results therefore suggested the existence of biological constraints over expression accumulation.

### Lineage tree-based analysis revealed a biological boundary of expression

We hypothesized that an expression boundary for individual genes could be one potential mechanism of noise constraint and tested this hypothesis with a simple geometric model for the gene expression dynamics (**Figure 3A**. See **Methods**). Briefly, the transcriptome of each cell is viewed as a particle with random motion within the *N*-dimensional (*N* is the number of genes, and in this case the 710 genes listed in **Table. S7**) expression space. Under this model, inheritance dictates that the transcriptome of all tips within a sub-tree should be limited by an *N*-sphere approximately centered at the root of the sub-tree, i.e. a limited range of expression variation (the circles in **Figure 3A**). If an expression boundary exists, a sub-tree whose root is closer to the boundary should display smaller range of expression variation than another sub-tree whose root is farther away from the boundary (e.g., gene 1 in **Figure 3A**). Note that although there should be both lower and upper boundaries for expression, we only focused on the upper boundary because the lower boundary is most likely undetectable given the overwhelming technical noise of scRNA-seq data towards the lower end of expression level.

**Figure 3.**
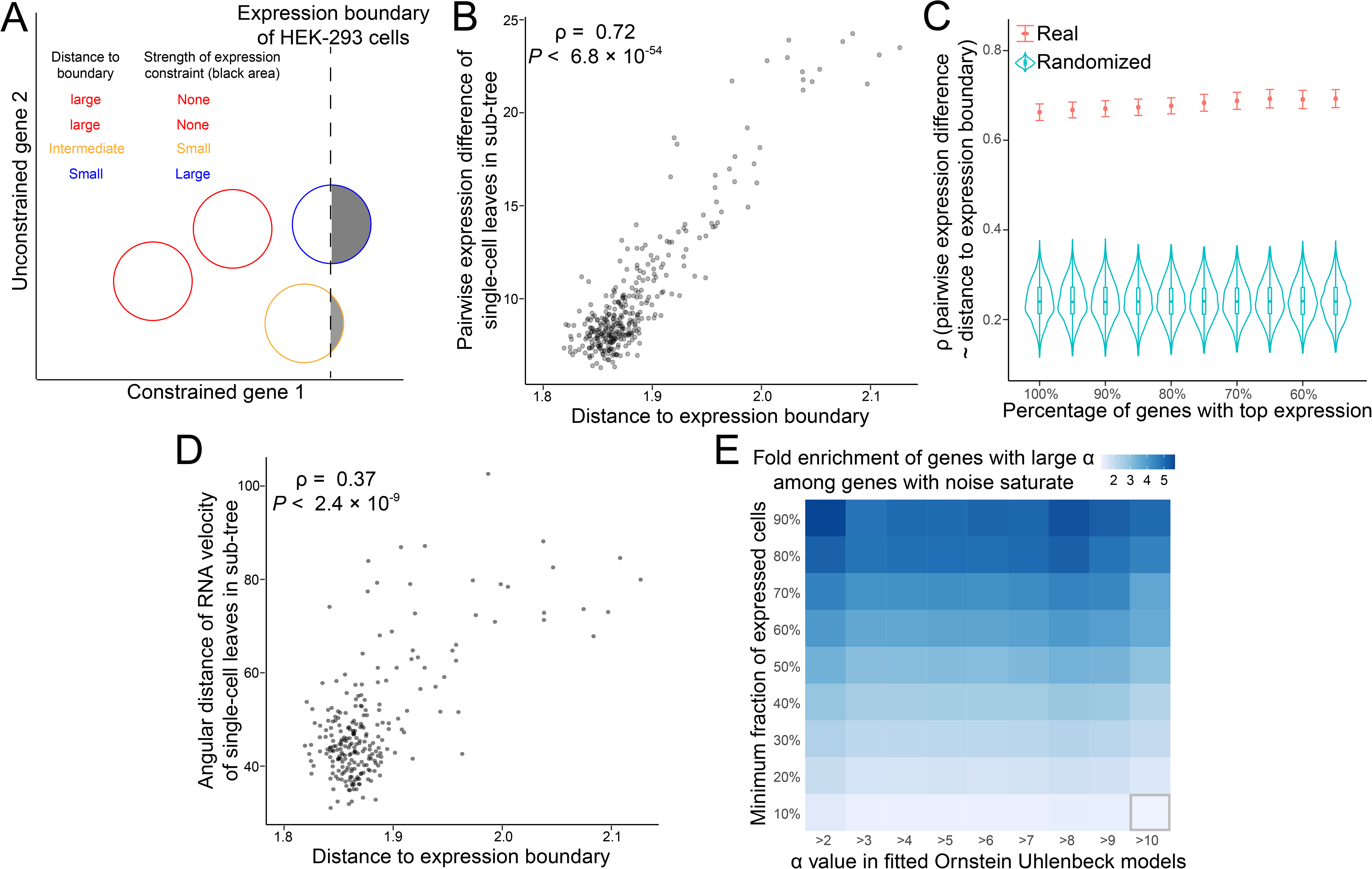
Evidence of biological boundaries for gene expression. **(A)** Schematic diagram of the geometric model of gene expression levels, indicating how it is used to detect the existence of a biological boundary of gene expression. A hypothetical gene whose expression is constrained by a boundary (*x* axis) and another hypothetical gene whose expression is not constrained by a boundary (*y* axis) are shown. Each circle represents the range of expression variation among the descendent cells of a single ancestor, which is presumably at the center of the circle. When the circle (and therefore the ancestor) is far away from the boundary (red circles), the full range of expression variation can be realized by the descendent cells. However, when the ancestor is closer to the boundary, such that some (the yellow circle) or a large (the blue circle) fraction of the circle is beyond the expression boundary (the gray fraction), the expression level of the descendent cells will be constrained and will appear less variable. **(B)** Consistent with the geometric model, a sub-tree that is farther away from the expression boundary (*x* axis) tends to show greater variation among the single-cell leaves (*y* axis). Each dot represents a sub-tree, and Spearman’s rank correlation coefficient is indicated along with the corresponding *P* value. **(C)** The positive correlations observed in B increased when a certain fraction (*x* axis) of highly expressed genes was used. The error bar represents the standard error estimated from 1,000 bootstrapping replicates for the genes. **(D)** For each real sub-tree, each single-cell leaf was replaced by other randomly selected single-cell leaves. These “fake” sub-trees constituted a “mock” dataset and were analyzed in a similar manner to panel c. The results from 1,000 such mock datasets are shown as violin plots representing the expected correlation, which is much lower than that observed from the real tree (as shown in c). **(E)** Ornstein Uhlenbeck model was fitted for each gene to detect the strength of mean-reverting tendency (_α_) of its expression level. We found that the 492 genes with noise saturation (**Table S7**) are significantly enriched of genes with large _α_ value, regardless the background list of genes defined by minimum fraction of expressed cells (*y* axis), or the threshold for the _α_ value (*x* axis). The fold enrichment is represented by the color of each tile, as indicated by the scale bar on top. A gray box around the tile indicates that the enrichment is not significant by hypergeometric test.

We inferred the upper boundary of expression of a gene by its maximal observed expression within the STADT data, and the transcriptome of the root of a sub-tree by hierarchically averaging the transcriptomes of its tips (See **Methods**). The difference between a gene’s expression in the root of a sub-tree and the upper boundary of the gene was calculated and these differences for all genes were geometrically averaged to yield the distance of the root to the expression boundary. We found that roots with smaller distance to the expression boundary tend to give rise to sub-trees with significantly less noisy tips (**Figure 3B**). Comparison with other published scRNA-seq data suggested that the inferred expression boundaries are unlikely physical, but instead most likely biological boundaries (**Figure S9**). In addition, limiting the above analysis to highly expressed genes, whose expression measurements are likely more accurate, strengthened the correlation (**Figure 3C**). More importantly, this pattern of less-noisy leaves in sub-trees closer to the expression boundary disappeared when we randomly rearranged the cells onto different leaves of the lineage tree (**Figure 3D**). In other words, the pattern of biological expression boundary is dependent on the real lineage relationship between the cells. To further support the existence of constraint on expression noise, we fitted the lineage tree and the expression level of each gene with Ornstein-Uhlenbeck (OU) model, which is commonly used in phylogenetic studies to infer mean-reverting tendencies of quantitative traits ^32^. As a result, we noticed that the 492 genes with significant noise saturation were enriched with genes with strong mean-reverting tendencies regardless various cutoffs for expression breadth among single cells of strength of mean-reverting tendency (**Figure 3E**). Collectively, our results suggested expression levels were individually constrained, such that expression noise is limited.

### Expression co-fluctuation due to transcriptional regulation and/or asymmetric division also constrains expression noise

The constraints on expression noise described above were one-dimensional, i.e., expression boundary for each gene is independent of other genes. However, another type of noise constraint, co-fluctuation between genes (**Figure 4A**), is also possible. Importantly, the lineage tree provided us with a unique advantage in resolving two possible mechanisms underlying co-fluctuation: transcript co-segregation in asymmetric division (**Figure 4B**) and transcriptional regulation (**Figure 4C**). Here, co-fluctuation caused by transcript co-segregation refers to the correlated biased distribution of transcripts between two sister cells due to molecular packaging (e.g. transcripts in one mitochondrion) which consequently become roots of two sub-trees that are highly divergent in terms of transcript abundance. The leaves of these two sub-trees may therefore exhibit expression levels clustered around their respective roots, thereby displaying signals of co-fluctuation between genes (**Figure 4B**). On the other hand, gene pairs showing co-fluctuation maintained by transcriptional regulation likely show strong correlations even during individual cell cycles (**Figure 4C**). This type of co-fluctuations is more likely regulated by extrinsic noise ^33^, such as genes regulated by the same transcription factor, or genes that are sensitive to changes in their relative dosage ratio (e.g. members of the same protein complex ^34^).

**Figure 4.**
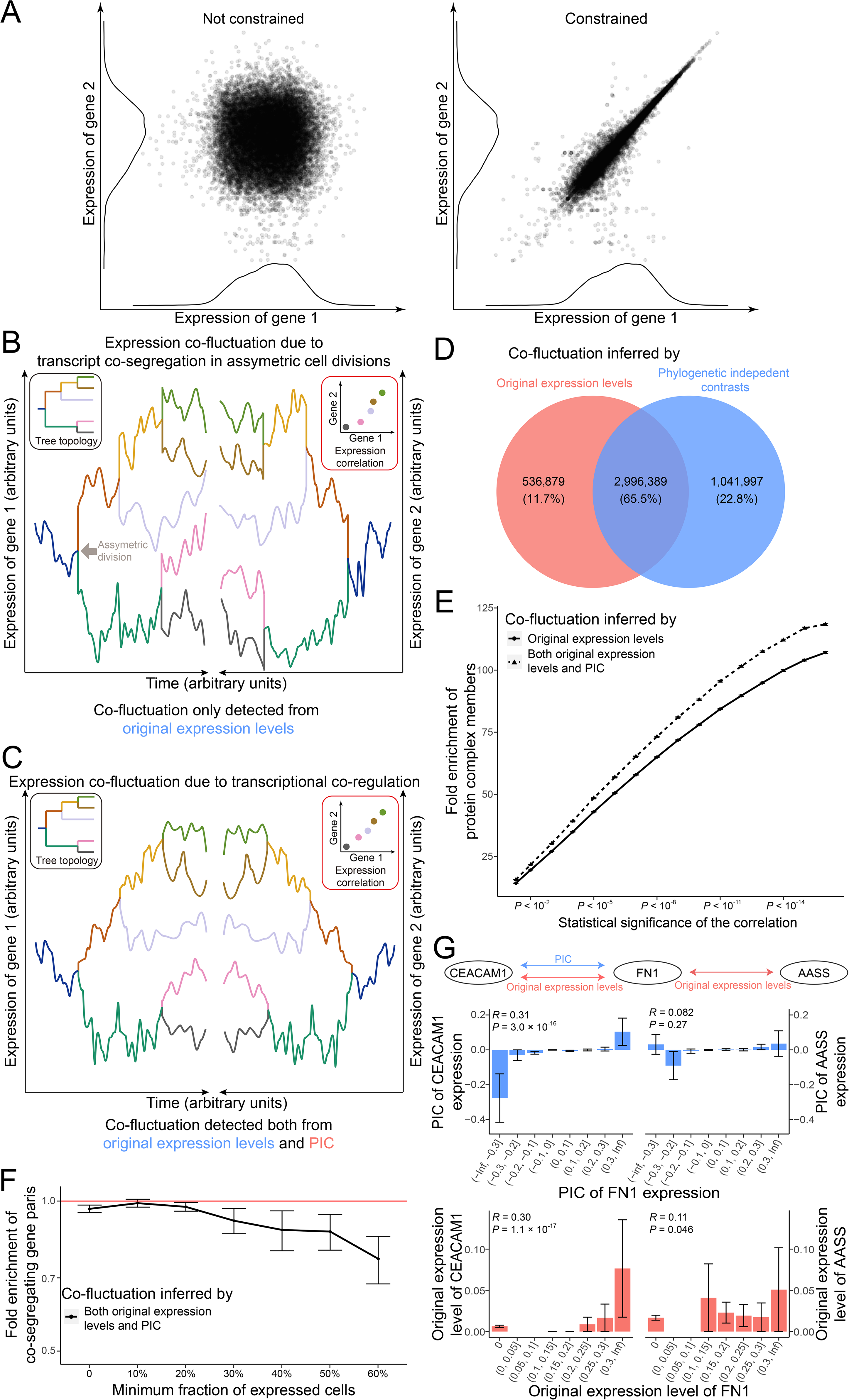
Co-fluctuation among genes also constrains gene expression noise. **(A)** Schematic diagram showing the expression levels of two hypothetical genes. On the left, the random fluctuation (noise) of the expression of the two genes is not correlated and is therefore unconstrained. On the right, the distribution of the expression of either gene remains individually consistent (histogram on the *x* and *y* axes, compared to those of the left panel), but the expression noise becomes correlated (i.e. co-fluctuation). The expression noise on the right is therefore more constrained than that on the left. (**B** and **C**) Two possible mechanisms mediating co-fluctuation, as demonstrated by the temporal (*x* axis) fluctuation of the expression levels (*y* axis) of two genes (one on the left and the other on the right). The lineage tree of the individual cells is indicated in the top left inset, with each branch color coded accordingly. The resulting pattern of co-fluctuation between the two genes among the five leaves of the tree is shown in the top right inset (**B**) Asymmetric cell division might have led to expression differences (and therefore noise) among sister cells, which are then inherited by descendent cells, resulting in expression co-fluctuation due to the shared history of ontogenetically closely related cells (also see Figure. S9). (**C**) On the other hand, transcriptional regulation could have maintained the dosage equality between the genes, resulting in expression co-fluctuation within individual cell cycles. (**D**) We used Pearson’s correlation coefficient of original expression level and phylogenetic independent contrast (PIC) analysis to assess expression co-fluctuation. The Venn diagram for gene pairs identified as significantly correlated by both or either of the two tests is shown, with the number of gene pairs indicated. (**E**) Fold enrichment (*y* axis) of gene pairs that are members of the same protein complex, which are assumed as sensitive to relative dosage change, are shown for co-fluctuating gene pairs inferred by different methods. The fold enrichment, as well as the enhancement by PIC, increases as we tighten up the statistical criteria for the correlation (*x* axis). Error bars indicate the standard deviation, estimated by bootstrapping gene pairs with indicated level of statistical significance 100□times. (**F**) The gene pairs inferred as co-fluctuating by both PIC and original expression levels are depleted (*x* axis) of co-segregating gene pairs relative to that inferred by original expression levels. The depletion becomes stronger (*y* axis) for genes expressed in more cells. Error bars indicate the standard deviation, estimated by bootstrapping genes within each cutoff 100□times. (**G**) On the left, two co-regulating genes show significant co-fluctuation inferred by both PIC and original expression levels. On the right, two co-segregating genes show significant correlation inferred by original expression levels but not by PIC. The coefficients of determination (*R*) between all gene pairs are showed.

To identify co-fluctuating gene pairs, we first simply calculated the Pearson’s correlation coefficients of the original expression levels for each pair among the 18,407 human genes with non-zero expression in our scRNA-seq dataset and identified > 3.5 million significantly co-fluctuating gene pairs (**Figure 4D**, left circle). To identify those pairs whose co-fluctuation was constrained by transcriptional regulation, we resorted to the method of phylogenetically independent contrast (PIC) analysis. PIC is required here as correlations directly calculated using the tips of the lineage suffer from statistical dependency due to shared ancestry, as is commonly shown in studies for correlated evolution of quantitative phenotypic traits^35^. Over 4 million pairs of genes (**Figure 4D**, right circle) were thus found to be likely co-fluctuating due to transcriptional regulation, over 74.2% of which were significant according to co-fluctuation inferred by the original expression levels (**Figure 4D**, overlap between the two circles). We noticed that gene pairs with significant co-fluctuations inferred by original expression levels are enriched with members of the same protein complex, which are sensitive to relative dosage changes ^34^ (**Figure 4E**, solid line). Such fold enrichment is, intriguingly, further increased if significant co-fluctuation inferred by PIC is also required (**Figure 4E**, dotted line). More importantly, this enrichment increment introduced by PIC (and therefore the lineage tree) became more dramatic if we tighten up the statistical criteria (smaller *P* value) for co-fluctuation (**Figure4E**).

Additionally, PIC should in theory capable of discriminating co-fluctuation due to transcript co-segregation. We tested this notion by detecting enrichment of gene pairs known to have their mRNAs colocalized on outer mitochondrion membrane ^36^. As a result, we found that co-fluctuating gene pairs inferred by both PIC and original expression level are depleted of colocalized mRNAs relative to co-fluctuating gene pairs inferred by only original expression level (**Figure 4F**). A closer look at the data revealed further some specific cases with apparent biological relevance. For example, two different adhesions, fibronectin 1 (FN1) and carcinoembryonic antigen-related cell adhesion molecule 1 (CEACAM1) are known to be co-regulated by Nuclear Factor Kappa B Subunit 1 (NFKB1) ^37–39^, and participated in the proliferation, migration, and invasion of cancer cells ^39–42^. The strong functional correlation of these two genes is consistent with the strong co-fluctuation inferred by either PIC or original expression level (**Figure 4G**, left). On the contrary, mRNA of FN1 and alpha-aminoadipic semialdehyde synthase (AASS) are co-localized on the outer mitochondrial membrane ^36^, and may be co-segregated during cell division. They appeared co-fluctuating by correlation of original expression levels, but not by PIC (**Figure 4G**, right).

## Discussion

To investigate the accumulation of expression noise during clonal expansion, we developed STADT, a novel technique combining scRNA-seq and genome editing, to simultaneously determine the single-cell transcriptome and high-density lineage tree of a single-cell clone consisting of ∼5,000 HEK-293 cells. Our dataset revealed rapid saturation of expression noise, biological boundaries of gene expression, and pervasive co-fluctuation between different genes, all of which point towards a highly constrained regulatory landscape of gene expression noise. To the best of our knowledge, this is the first comprehensive study of the regulation of gene expression noise in the context of clonal expansion and therefore substantially deepens our understanding of how the stochasticity of gene expression is constrained in the context of ongoing cell divisions.

A few limitations of our analyses are worth discussing. First, the estimation of divergence times between cell pairs relies heavily on the accuracy of the lineage tree reconstructed by STADT. The divergence time could be underestimated due to inevitable cell losses during the growth of the single-cell clone or library preparation using 10x Chromium technology or might be overestimated due to false editing events caused by STADT barcode sequencing errors. To minimize the impact of these biases, we optimized the STADT protocol to retain as many cells as possible, used PacBio HiFi reads to ensure highly accurate readout of STADT barcodes, and chose an estimate of the divergence time that was robust to underestimation due to cell losses. More importantly, the decrease in the share of ancestral STADT allele transcripts and the increase in expression differences with increasing divergence time strongly suggested that the hierarchical relationship/kinship between the cells was generally accurate. Second, the single-cell transcriptome quantified with 10x Genomics Chromium technology is subject to high technological noise. For example, it was previously shown that droplet-based scRNA-seq shows a CV of at least 0.5 for the counts per million (CPM) values of the majority of genes ^43^. Nevertheless, the patterns revealed in our analyses as well as by other researchers^44^ seem to suggest that such data might still capture the major trends of expression noise, such as co-fluctuation between genes ^44^. However, our results might have been undermined by such technological noise, suggesting the existence of even stronger biological signals underlying our observations. Third, we did not discriminate extrinsic noise, which arises from cell state difference, and intrinsic noise, which arises from the stochasticity during gene expression of cells of the same state. We, however, suspected that the noise constraints revealed by STADT data are mostly for extrinsic noise, as both expression boundary and co-fluctuation likely require extrinsic regulatory factors. It will be interesting to further disentangle intrinsic and extrinsic noise using methods such as allele-specific scRNA-seq data ^30^.

STADT represents a technological effort in a novel (to our knowledge) direction of scGESTALT-like assays for the reconstruction of cell lineage trees, in which there is more emphasis on the sampling density in terms of single cells, instead of the breadth of sampling ^16, 17^. Further possible modifications of STADT for higher sampling densities include longer STADT barcodes for a larger lineage recording capacity, cell cycle-coupled editing for a better resolution of division events, and the replacement of the CRISPR/Cas9 system (which causes DNA double-strand breaks and, thus, DNA damage-induced cell death) with other genomic editing systems. Nevertheless, our current study demonstrated the unique advantage of such dense lineage trees in analyzing gene expression noise. For example, the noise saturation occurring at a divergence time of 3 ∼ 4 cell divisions would have been missed by lineage trees with a lower density because most division events would be untraceable due to the high number of cell losses. We would also like to note that in terms of approximating divergence time between individual cells, high-density sampling for cells might be a promising alternative to statistically inference based on assumed editing efficiency of Cas9 on the lineage barcode, because the editing efficiency of Cas9 is much more variable than the distribution of loss cells, especially when the single cells are sampled with sufficiently high density. Additionally, we conducted the first transcriptome-wide analyses of expression noise in single cells controlling their kinship, an underappreciated factor in previous scRNA-seq-based studies of expression noise. The dense sampling of single cells by STADT should further increase the statistical power for detecting co-fluctuation between genes ^45^.

Our analyses of gene expression noise provided insights for more fundamental questions in cellular biology. For example, how does a growing population of cells maintain identity/functional types? (**Figure 5**) The aforementioned technological limitations notwithstanding, our results provide a clear answer: gene expression differences among cells are biologically constrained at levels ranging from individual genes (expression boundary) to interactions among genes within the same cell (co-fluctuation), to populations of cells (noise saturation). We also want to emphasize that the evidence presented here for the mechanisms of homeostatic maintenance of transcriptome critically rely on the knowledge of the (dense) lineage tree, which provides a unique advantage in supplement to static snapshots of either bulk or single-cell RNA-seq results. These results also point to potential applications, such as assessing the differentiation of cells and therefore the objective definition of cell types ^46–48^ according to an excessive accumulation rate of expression differences among cells, and suggest candidate strategies for cell fate manipulation based on an individual gene’s expression boundaries or co-fluctuation among genes. Altogether, our results substantially enrich our understanding of the regulation of expression noise and the biological basis of homeostatic maintenance of cell identity. It will be interesting in the future applying STADT to assess the differentiating and non-differentiating lineage trees of the same type of cells, or to compare the lineage tree underlying the expression geometry among normal cells with that among cancerous cell ^49^.

**Figure 5.**
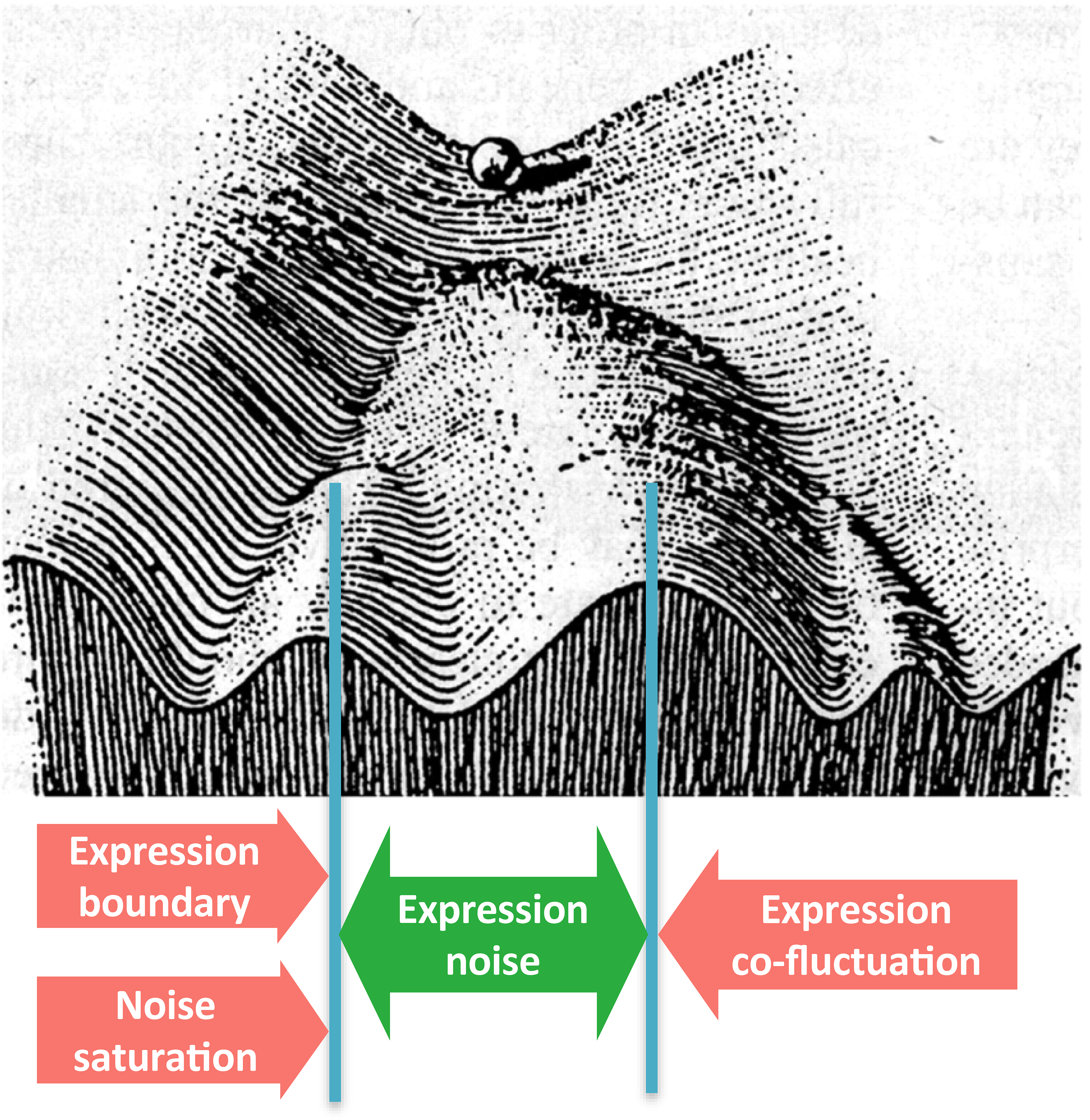
Role of expression noise and its constraint in the context of Waddington’s landscape model. In Waddington’s seminal work^23^, he metaphorically described the developmental process of a biological system (e.g. a cell) as a marble rolling down a slope with ridges and depressions, where the depressions at the base of the slope represent stable differentiated states (cell identity), and the ridges the (epigenetic) barriers between these states. Under this model, expression noise resembles an inherent tendency to cross the ridge (the green two way arrow) and therefore conversion from one cell type to another. On the contrary, the constraints of expression noise identified in this study resemble forces keeping the marble within the depression (the red arrows) and therefore regulatory mechanisms to maintain cell identity and homeostasis.

## Methods

### Design and construction of the STADT barcode and sgRNAs

To design the STADT barcode and corresponding sgRNA, we first generated > 1 million randomized 20 bp sequences as candidate sgRNAs, among which the sgRNAs with no more than three substitutions relative to any fragment of the human genome were removed for further analysis. From the remaining sgRNA candidates, we selected the spacer sequence 5’-TATTCGCGACGGTTCGTACG-3’, which was designated sgRNA1. Then, 13 protospacer sequences were designed based on sgRNA1 according to the following criteria: (i) each protospacer contained 2-3 mismatches with sgRNA1, (ii) there was no recurrence of any sequence of 9 bp or longer, and (iii) consecutive repeats of the same nucleotide for more than 2 bp were completely absent. The 13 protospacers along with protospacer-adjacent motif (PAM) sequences designed following a previous strategy ^50^ were concatenated in an order of decreasing cutting frequency determination (CFD) scores ^51^ to obtain the STADT barcode. To further ensure that the lineage recording capacity of the STADT barcode was fully utilized, we designed sgRNA2/3/4, which targeted the 9^th^/12^th^/13^th^ protospacer sequences (rarely edited sequences in preliminary experiments) with 100% identity but showed small CFD scores (< 0.55) with other protospacers. The designed STADT barcode and sgRNA sequences are listed in **Table S1**.

To obtain a lineage tree with a high density, it is crucial that during single-cell library preparation with 10x Chromium technology, the STADT barcode can be successfully reverse transcribed. We therefore applied multiple approaches to enhance the capture rate of the STADT barcode by 10x Chromium technology, including (i) using the strong promoter EF1_α_, (ii) concatenating the STADT barcode with the 3’ end of the gene encoding EGFP so that green fluorescence could be used as a marker for single-cell colonies with appropriate expression of the STADT barcode, and (iii) adding a 20 nt poly-dA (A_20_) sequence at the 3’ end of the STADT barcode to facilitate capture by the poly-dT RT primer on 10x gel beads. Along with other accessory elements, the final STADT barcode vector (pLV-EF1A>EGFP:T2A:Bsd:V1. VectorBuilder, no. VB170906-1089zqd) was constructed by using Gateway technology ^52, 53^ on the backbone of pLV-3d. After the transfection of the STADT barcode vector into HEK-293 cells (see below), we applied strong selection by blasticidin (15 µg/ml) for 12 days to ensure a high expression level of the STADT barcode. For the sgRNAs, two DNA cassettes, U6>sgRNA1-U6>sgRNA2 and U6>sgRNA3-U6>sgRNA4, were synthesized and subsequently combined by Golden Gate ligation in the sgRNA vector pLV-U6>sgRNA1-U6>sgRNA2-U6>sgRNA3-U6>sgRNA4 (VectorBuilder, no. VB180515-1178wxn).

### Construction of the HEK-293 cell line for lineage tree reconstruction

The two lentiviral particles of the barcode and sgRNA vectors were individually transfected into HEK-293FT cells (Cyagen, no. HEKFT-30001) with lentivirus packaging plasmids using Lipofectamine 2000 (ThermoFisher, no. 11668027). To obtain the viral particles, the supernatant was harvested three days post-transfection, in which cellular debris was removed by centrifugation at 3000 rpm for 15 min and filtering with a 0.45 µm disposable needle filter (Millipore, no. SLHP033RB), followed by centrifugation at 50,000 ×g for 90 minutes. To gauge the viral titration, the viral particle-containing supernatant was diluted to various concentrations, which were then used to transduce a second batch of HEK-293FT cells. The genomic DNA (gDNA) of these HEK-293FT cells was isolated using DNeasy Blood & Tissue Kits (Qiagen, no. 69504). The gDNA was further analyzed by qPCR targeting the barcode and the endogenous gene *BMP2* using the QuantiNova SYBR Green PCR kit (Qiagen, no. 208057-500T) so that the viral titration could be determined.

The inducibility of *Cas9* in the Inducible Cas9 Expression 293 Cell Line (abm, no. T3352) was confirmed by Western blotting in cells cultured in the presence of different concentrations of cumate (abm, no. CH065). These cells were then transduced with lentiviral particles containing STADT barcodes at a low MOI (0.1), followed by selection with 15 µg/ml blasticidin (InvivoGen, no. ant-bl-1) for 12 days, giving rise to HEK-293 cells harboring a single copy of the STADT barcode. The sgRNA lentiviral particles were transduced at different MOIs (10, 20, and 30), and positive cells were selected with 400 µg/ml hygromycin (InvivoGen, no. ant-hg-1) for 5 days. PCR or RT-qPCR was used to detect the expression levels of Cas9, the STADT barcode and sgRNA1 using the primers listed in **Table S2**. The constructed HEK-293 cells containing inducible Cas9 and the STADT barcode/sgRNA are hereafter referred to as STADT-HEK-293 cells.

### Gauging the editing efficiency of STADT barcode

Approximately 10^6^ STADT-HEK-293 cells were grown in 10% serum media containing 30 µg/ml cumate in a 6-well dish. After three days of culture, cells were harvested for gDNA extraction using DNeasy Blood & Tissue Kits (Qiagen, no. 69504). Alleles of the STADT barcode were amplified from gDNA using Phanta Max Super-Fidelity DNA Polymerase (Vazyme, no. P505) with the gDNA-V1f and gDNA-V1r primers (**Table S2**), and cloned into the CE Entry Vector (Vazyme, no. C114). Recombinant plasmids were obtained by blue-white selection, and 20 randomly selected clones were subjected to Sanger sequencing to assess the editing efficiency.

To gauge the editing efficiency of the STADT barcodes in single-cell clones, 288 single cells were first manually selected by micromanipulation from the log-phase population of STADT-HEK-293 cells constructed by using three different titrations of sgRNA lentiviral particles (pLV-sgRNA, MOI = 10, 20, and 30). These single cells were then grown in three 96-well plates for 10 days, during which 10 µl of CloneR^TM^ (STEMCELL, 05888) was added to the medium of each colony on the 0^th^ and 1^st^ days, while 5 µg/ml and 30 µg/ml cumate were added to the media on the 0^th^, 2^nd^, 4^th^, 6^th^, and 8^th^ days. The colonies were visually inspected and categorized as follows: normal (type I), disassociated/substantial death (type II) or not-growing/excessive death (type III). Thirty type I single-cell colonies with different expression levels of sgRNA and Cas9 were chosen. Then approximately 50 ng of gDNA was harvested from each colony using the QIAamp DNA Micro Kit (Qiagen, no. 56304). The gDNA was PCR amplified in a 50 µl reaction targeting the STADT barcode, whose editing status was further confirmed by Sanger sequencing (using 13 µl of the product), TA cloning (1 µl of the product) or Illumina HiSeq-PE250 sequencing (36 µl of the product) as follows.

Based on the Sanger sequencing data, we found that ∼93% of the STADT alleles from these single-cell clones were found to be edited (**Figure S4A**). Individual STADT alleles determined by TA cloning and subsequent Sanger sequencing showed that colonies with high copy numbers of sgRNA in the genome (transformation with MOI = 20 or 30) that were grown in the presence of high concentrations of cumate (5 or 30 µg/ml) displayed a reasonably high diversity of alleles (**Figure S5B-G**).

The sufficient editing efficiency and diversity of STADT was further corroborated by the sequencing data from Illumina HiSeq-PE250. Briefly, the clean data of each of the eight sample sequenced by HiSeq-PE250 were trimmed with fqtrim (https://ccb.jhu.edu/software/fqtrim/) using the default parameters. Reads 1 and 2 were spliced with flash^54^ using 30 bp of overlapping sequence and 2% mismatches. Nonspecifically amplified sequences generated during PCR were removed by Bowtie2 ^55^ with alignment to the human reference genome using default parameters. Falsely amplified sequences were further removed by filtering according to the two primers gDNA-V1f and gDNA-V1r (**Table S2**) allowing 2 mismatches. After these two rounds of filtering, most of the remaining sequences amplified from the STADT barcode were retained. Finally, the sequences of the STADT barcode were aligned with the reference sequence by MUSCLE^56^ using the default parameters, and the editing events in each sequence were called according to the methods proposed in the GESTALT paper^50^ (**Figure S5A-H**). For the sample (**Figure S4H**) with the highest sgRNA copy number (MOI = 30) and strongest cumate induction (30 µg/ml), ∼ 82.9% of the deletion events revealed by HiSeq were shorter than 20 bp (**Figure S4I**), and majority of the STADT alleles contained no inter-site deletions (**Figure S4J**). More importantly, the frequency of editing events suggested that editing started as early as the two-cell stage of the colony (**Figure S4K**), and long deletion events were absent until the 16-cell stage (**Figure S4L**). These features suggested that STADT utilized the lineage recording capacity of the barcodes better than previously reported scGESTALT-like techniques.

### Preparation of single cells for 10x Genomics scRNA-seq

To assess the feasibility of clonal expansion starting from a single cell, we prepared 384 single-cell clones of the STADT-HEK-293 cells constructed with the sgRNA lentiviral particles at an MOI = 30. Similar to the cultivation procedure described above, we added 10 µl of CloneR^TM^ and 30 µg/ml of cumate to the medium of each colony at different time points (see above) and allowed the colonies to grow for 10 days. The colonies were again visually inspected and categorized as follows: normal (type I, 43 colonies), disassociated/substantial death (type II, 162 colonies) or not-growing/excessive death (type III, 179 colonies) (**Figure S3A**). The results suggested that despite a relatively low success rate, the generation of a STADT-HEK-293 population seeded from a single cell is feasible.

We then used an optimized protocol for single-cell digestion to minimize cell loss, which is detailed as follows. First, the culture medium was removed by gentle suction, and morphologically normal colonies (type I colonies remaining after suction, according to visual inspection under an optimal microscope) were retained. Second, without washing the single-cell colonies with any solution, we directly added 100 µl of ACCUTASE (StemPro, no. A1110501) to digest the colonies at 37 °C for 20 ∼ 30 minutes and continuously examined the status of the single-cell clones after 20 min. Third, the colonies were disassociated into single cells via gentle, repetitive pipetting, after which 900 µl of PBS was added to each colony to terminate the digestion. Finally, each 1 ml single-cell suspension was centrifuged at 500 ×g for 10 min. The supernatant was removed by gentle suction and discarded, and the remaining single cells were resuspended in 20 µl of PBS containing 5% BSA. Altogether, we obtained single-cell suspensions for seven samples (among 15 colonies at the beginning of digestion). We used two of these samples to determine the fraction of living single cells by trypan blue staining and cell counting with Countstar BioMed IM1200, which indicated 95% (among ∼ 3300 cells) and 93% (∼ 3800 cells) living cells, respectively (one sample in **Figure S3C**).

As a final batch of single-cell colonies to be used for scRNA-seq, we prepared 576 single-cell clones of STADT-HEK-293 cells constructed with sgRNA lentiviral particles at an MOI = 30, and we cultured the clones similarly for 10 days (see above). We discarded type II and type III colonies and colonies with weak GFP fluorescence (i.e., STADT barcode expression), resulting in 15 remaining colonies that were subjected to further digestion. The number of cells within each colony was determined from the ratio between the area of the colony and that of a typical single cell (**Figure S6A and B**).

### Transcriptome and lineage barcode sequencing

The other five single-cell suspension samples were loaded onto a GemCode Single-Cell Instrument (10x Genomics, Pleasanton, CA, USA) in their entirety to obtain the GEM-RT product following the Single Cell 3’ Reagent Kit v2 User Guide. Half of the GEM-RT product was directly amplified without being fragmented, targeting the STADT barcode (Sc-barcode-f and Sc-barcode-r listed in **Table S2**), and the product was later subjected to TA cloning followed by Sanger sequencing of ∼ 20 STADT alleles to obtain a rough estimation of the diversity of the STADT barcode. The two samples showing the highest diversity of STADT barcodes were chosen and sequenced with PacBio Sequel technology via standard library preparation and sequencing protocols. The remaining half of the GEM-RT product of each of these two chosen samples was cleaned, fragmented, amplified, and used to prepare a sequencing library with an average fragment size of 450 bp. Library sequencing was finally performed on the Illumina NovaSeq platform following a standard 10x scRNA-seq protocol.

### Analysis of scRNA-seq

For the STADT-HEK-293 samples assayed in this study, raw scRNA-seq reads were obtained from the Illumina Nova-seq platform. Mapping and quantification were performed with the reference GRCh38 using Cell Ranger ^57^ according to the official 10x Genomics protocols, which resulted in a count table (of unimolecular identifiers, or UMI) for each gene in each droplet/cell. Cells with < 200 detected genes or a > 9% mitochondrial content (fraction of reads derived from mitochondrial genes) were removed. Expression profiles were obtained for > 58% of the single cells loaded into the GemCode Single-Cell Instrument (**Figure S6A and B**). Expression data were normalized for sequencing depth using the NormalizeData function in the Seurat R package ^58^. For the analysis of the physical or biological boundaries of expression, raw expression matrixes of STADT-HEK-293 cells and multiple other scRNA-seq datasets ^16, 17^ generated with Cell Ranger were combined and normalized according to sequencing depth by Seurat. Under the geometric model of expression levels, the boundary of expression for each individual gene was estimated by using the maximal observed expression level among all single cells. Then, for each cell, the distances to the expression boundary for all genes were combined according to their geometric average (*x* axis of **Figure 3B**).

### Construction of lineage trees

To distinguish the heteroduplex DNA formed during PCR, all sub-reads from each zero-mode waveguide (ZMW) well on the PacBio Sequel platform were first categorized as belonging to the positive or negative strand according to the existence of 12 successive A or T bases, respectively, within the raw sequence (allowing three mismatches except in the leading three A/T bases). Consensus sequences were generated for both positive and negative strands by using the official CCS program ^59^ from PacBio, with a minimum of 3 passes/subreads per strand. The consensus sequences were then aligned to the human genome by using Bowtie2 ^55^ with default parameters, and the sequences with significant hits were removed because they represented nonspecific amplification with the STADT barcode primers. We further checked the existence of four STADT barcode primers (Sc-barcode-f, Sc-barcode-r, gDNA-V1f, and gDNA-V1r, in **Table S2**) within the consensus sequence, allowing two mismatches for each primer. Next, the cell barcode and UMI were extracted from the consensus sequence and were refined according to the cell barcode and UMI in the transcriptome data. That is, if the cell barcode and UMI from Sequel or NovaSeq differed by only one nucleotide, they were considered to be the same. Finally, the STADT barcode sequences were extracted from the remainder of consensus sequences between the fragments corresponding to primers gDNA-V1f and gDNA-V1r (**Table S2**). Detailed summary statistics of the read numbers throughout the analyses presented above are listed in **Table S3**.

To further correct the sequencing error on the PacBio Sequel Platform, the STADT barcodes with the same cell barcode and UMI were merged by multiple sequence alignment with MUSCLE ^56^, followed by selecting the nucleotide with the highest frequency at each site. Then, the STADT barcodes were aligned with the reference sequence by MUSCLE using the default parameters, and the editing events of each sequence (or STADT allele) were called according to a previous method employed by GESTALT ^50^. Then, for each STADT allele, the frequency was calculated as the total number of UMIs of the allele and its ancestral STADT allele. Here, the ancestral of a specific STADT allele was defined as any STADT allele in which the observed editing events were a subset of the editing events in the focal STADT allele; i.e., the focal STADT allele could be generated by further editing of the ancestral STADT allele (**Figure 1E**). Finally, the STADT allele of a cell was defined as the allele with the highest frequency, prioritizing the STADT alleles with more editing events if the frequencies were equal.

We sequenced 2863 (sample A) and 2294 (sample B) cells from the two scRNA-seq samples, among which >93% of the cells also had STADT barcodes recovered from PacBio Sequel. Finally, all recovered STADT barcodes of the two samples were reconstructed into multifurcating lineage trees using maximum likelihood implemented via the IQ-TREE LG model ^60^. To process our data with IQ-TREE, we replaced the default amino acid substitution model matrix in IQ-TREE with a matrix containing integers from 1 to 190, which represented the weights of editing events. Accordingly, all STADT alleles were encoded in a “sequence” of “amino acids”, with each “residue” in the sequence representing one editing event, and the ancestral and derived states of each “residue” were chosen according to the aforementioned substitution matrix, such that rare editing events received small weight scores and common editing events received large weight scores. Finally, we obtained lineage tree of which > 91% of the leaves were single cell. The detailed statistics during the above data processing steps are listed in **Table S4**. Programming codes for reproducing our analyses can be found on GitHub.

### Estimating divergence time between single cells

Under the assumption that the sub-trees were all unbiased samples of clonal expansion (**Figure S1**), a quantitative metric was required to put each sub-tree to the context of a certain stage of clonal expansion, such that the accumulation of expression noise could be shown alongside the progress of clonal expansion. An intuitive choice for such a metric might be the number of leaves (or cells in the leaves) of the sub-tree; however, this metric might be rendered inappropriate due to the nonnegligible fraction of cells that were not captured by the tree. Instead, we used the depth of the sub-tree, defined as the length (in units of the number of cell divisions) of the longest branch in the sub-tree (**Figure 2A**). Using the depth of sub-trees has several unique advantages. First, divergence time estimated by tree depths better suited our analyses in the context of cell divisions. Second, using tree depth provides estimates of the progress of clonal expansion that were more robust against the loss of cells compared to using the tree size (number of tips). This is because the impact of errors in depth estimates for small sub-trees should be weakened by the vast number of small sub-tree, and that for big sub-trees should be negligible as the depth should be correct with merely one intact branch (no cell loss on the branch). Third, increasing sampling density for cells, which is a major feature of STADT, should reduce errors in divergence estimates based on tree depths, but not those based on assumed editing efficiency of Cas9 on the lineage barcode. In the extreme case of 100% cell capture, depth-based estimates of divergence should contain no error. On the basis of the depth of the sub-tree, the divergence time between any pair of cells was then defined as the depth of the sub-tree rooted at the most recent common ancestor.

### Fitting the OU models

For each gene, its normalized expression levels in all individual single cell leaves, in combination with the overall lineage tree, were processed by the “ouch” package ^32^ in R to fit the OU model.

### Fold enrichment analysis

Take the enrichment of co-fluctuating gene pairs in protein complex as an example, we constructed a 2×2 contingency table:

|  | Members of the same<br>protein complex | Not members of the same<br>protein complex |
| --- | --- | --- |
| Expression correlated | *A* | *B* |
| Expression not correlated | *C* | *D* |

by categorizing each pair of genes (only those detected by scRNA-seq) into one of the four cells in the table on the basis of (i) whether the pair of genes showed correlated (by either Pearson’s correlation coefficient or PIC) expression and (ii) whether both genes in the pair were members of the same protein complex or co-localized on the outer mitochondrial membrane. The fold enrichment was calculated as (*A*/(*A*+*C*))/((*A*+*B*)/(*A*+*B*+*C*+*D*)). The fold enrichments of genes with strong mean-reverting tendency among the 492 genes with significant noise saturation were calculated similarly.

### Phylogenetic independent contrasts and co-fluctuation among genes

Phylogenetic independent contrasts (PIC) were calculated in R using normalized expression levels of individual genes and the lineage tree. Note that the linear models fitted to find correlation between PICs of different genes are required as going through the origin. But the same requirement did not apply to correlations for the original expression levels. The list of human protein complexes and their members were obtained from the CORUM database ^61^. To examine whether co-fluctuations inferred by PIC can eliminate co-segregating gene pairs, we also extracted the list of genes localized to the outer mitochondrial membrane (OMM) from previous study ^36^, which was defined as mRNAs with a four-fold enrichment in OMM APEX-seq over negative controls.

## Acknowledgments

We thank Xionglei He, Wenfeng Qian, Jianzhi Zhang, Mengyi Sun, Brian Metzger, Fabien Duveau, Zheng Hu, Zhengting Zou and Kehui Liu for their comments on the manuscript. This work was supported by the National Key R&D Program of China (grant number 2017YFA0103504 to X. C.) and the National Natural Science Foundation of China (grant numbers 31871320 and 81830103 to J.-R. Y., 31771406 to X. C., 32000401 to F. C.). All raw data from high-throughput sequencing have been deposited in NCBI BioProjects under accession number PRJNA715570 (to be released). Custom R and Python codes were used in data analysis and are available on GitHub (https://github.com/chenfengokha/noiseOnTree).

## Author contributions

Conceptualization, J.-R.Y.; Methodology, F.C., Z.L., X.Z., X.C. and J.-R.Y.; Software, F.C., Z.L.; Formal analysis, F.C. and Z.L.; Investigation, F.C., Z.L., X.Z., P.W., W.Y., X.C. and J.-R.Y., Resources, X.C. and J.-R.Y.; Data Curation, F.C. and Z.L.; Writing – Original Draft, F.C., Z.L. and J.-R.Y.; Writing – Review & Editing, F.C., X.C. and J.-R.Y.; Visualization, F.C. and Z.L.; Supervision, X.C. and J.-R.Y.; Project Administration, X.C. and J.-R.Y.; Funding Acquisition, F.C., X.C. and J.-R.Y.

## Supplementary Information

### Supplemental Figure Legends

**Figure S1.**
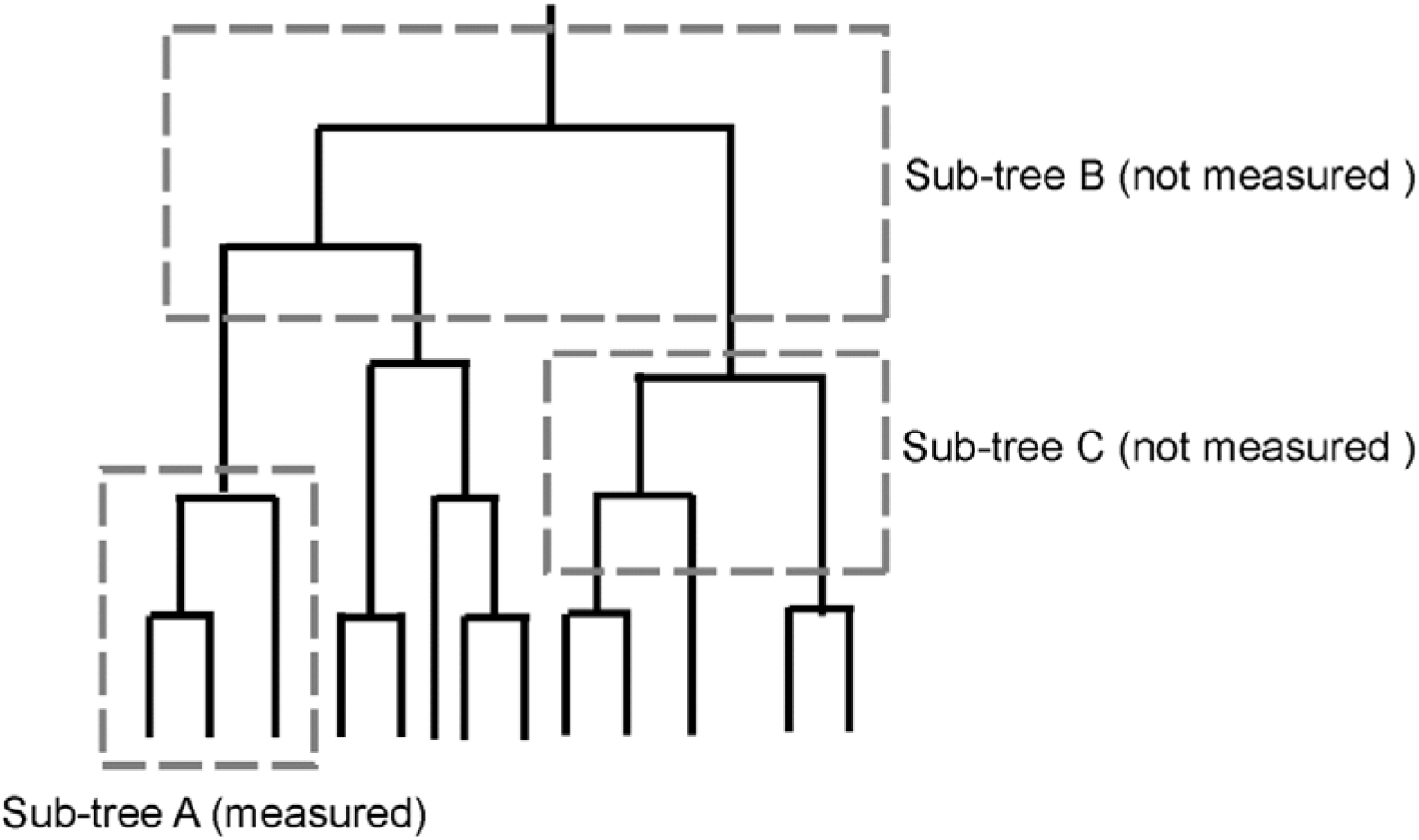
Schematic diagram for a basic assumption of the presented analyses. A random lineage tree is presented. Three gray boxes outline three different sub-trees. One of these sub-trees (sub-tree A) was directly measured, as the leaves of the sub-tree were assayed by scRNA-seq. Two other sub trees (sub-tree B and C) were not directly measured. Ideally, we would like to obtain the single-cell transcriptomes for all “internal” cells of the full tree, such as the leaves of sub-trees B and C, which is currently impossible because assessing the single-cell transcriptomes for the internal cells involves killing these cells. We therefore have to assume that sub-trees A, B and C are random samples of the same distribution. In other words, sub-tree A can be considered sub-tree B of an unrealized clonal expansion (and therefore its underlying full lineage tree) seeded by the root of sub-tree A. Similarly, sub-tree A can represent sub-tree C.

**Figure S2.**
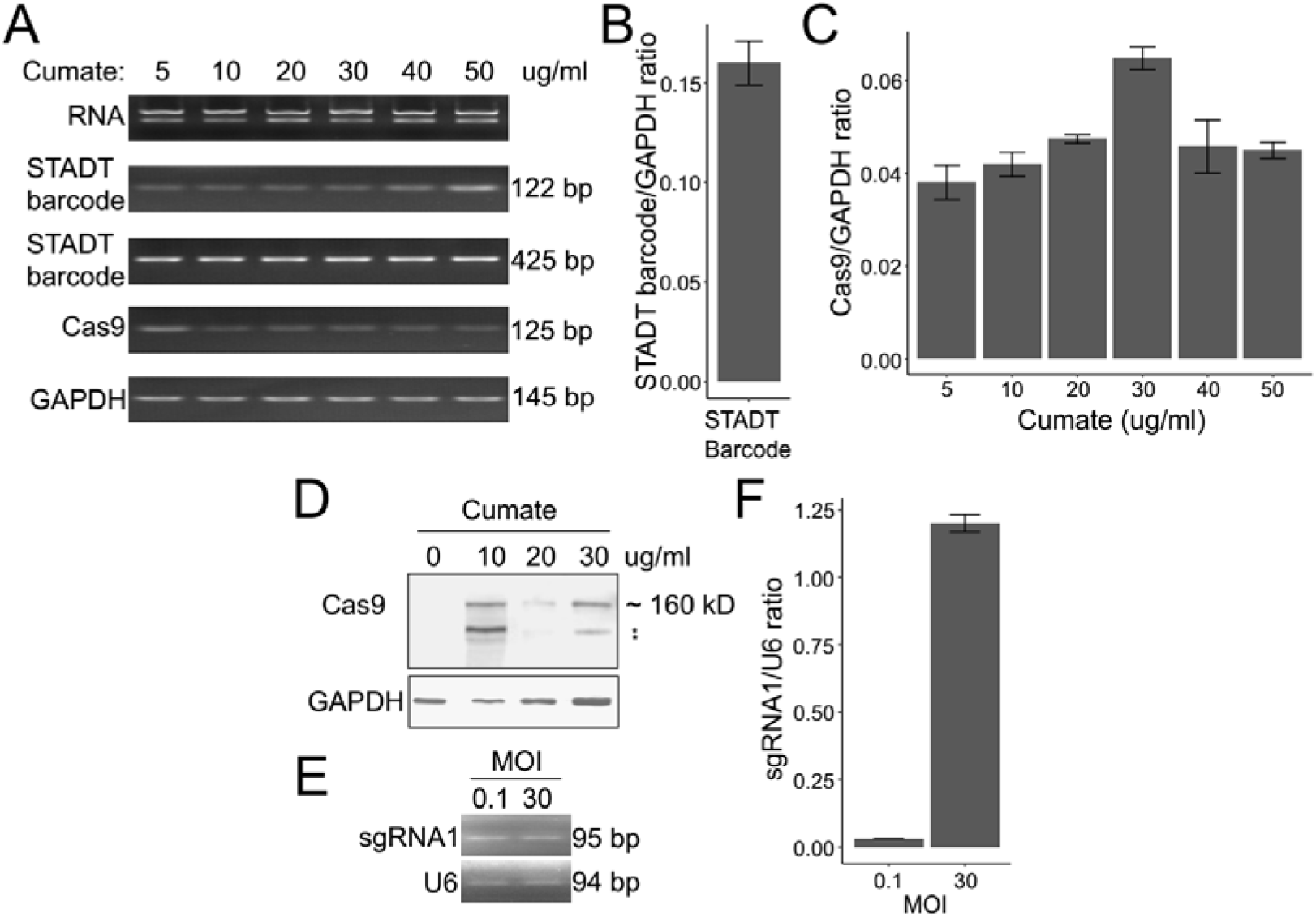
Validating expression levels of STADT components. (**A**) RT-PCR results for the RNA expression levels of the STADT barcode and Cas9 in cells growing in the presence of different concentrations of cumate. GAPDH was used as an internal control. The STADT barcode and Cas9 were amplified for 30 cycles, while GAPDH was amplified for 26 cycles. (**B**) RT-qPCR results for the expression levels of the STADT barcode relative to those of GAPDH. (**C**) RT-qPCR results for the expression levels of the Cas9 gene relative to those of GAPDH in cells cultured with different concentrations of cumate. (**D**) Western blot of the expression levels of Cas9 in cells cultured with different concentrations of cumate. GAPDH was used as an internal control. The small bands (*) may have resulted from the degradation of the Cas9 protein. (**E**) RT-PCR results showing the expression levels of sgRNA1 in the cells infected with different MOIs (0.1, 30) of the sgRNA lentivirus. U6 was used as an internal control. sgRNA1 was amplified for 35 cycles, while U6 was amplified for 30 cycles. (**F**) RT-qPCR result showing the expression level of sgRNA1 relative to that of U6 in the cell lines infected with different MOIs (0.1, 30) of the sgRNA lentivirus.

**Figure S3.**
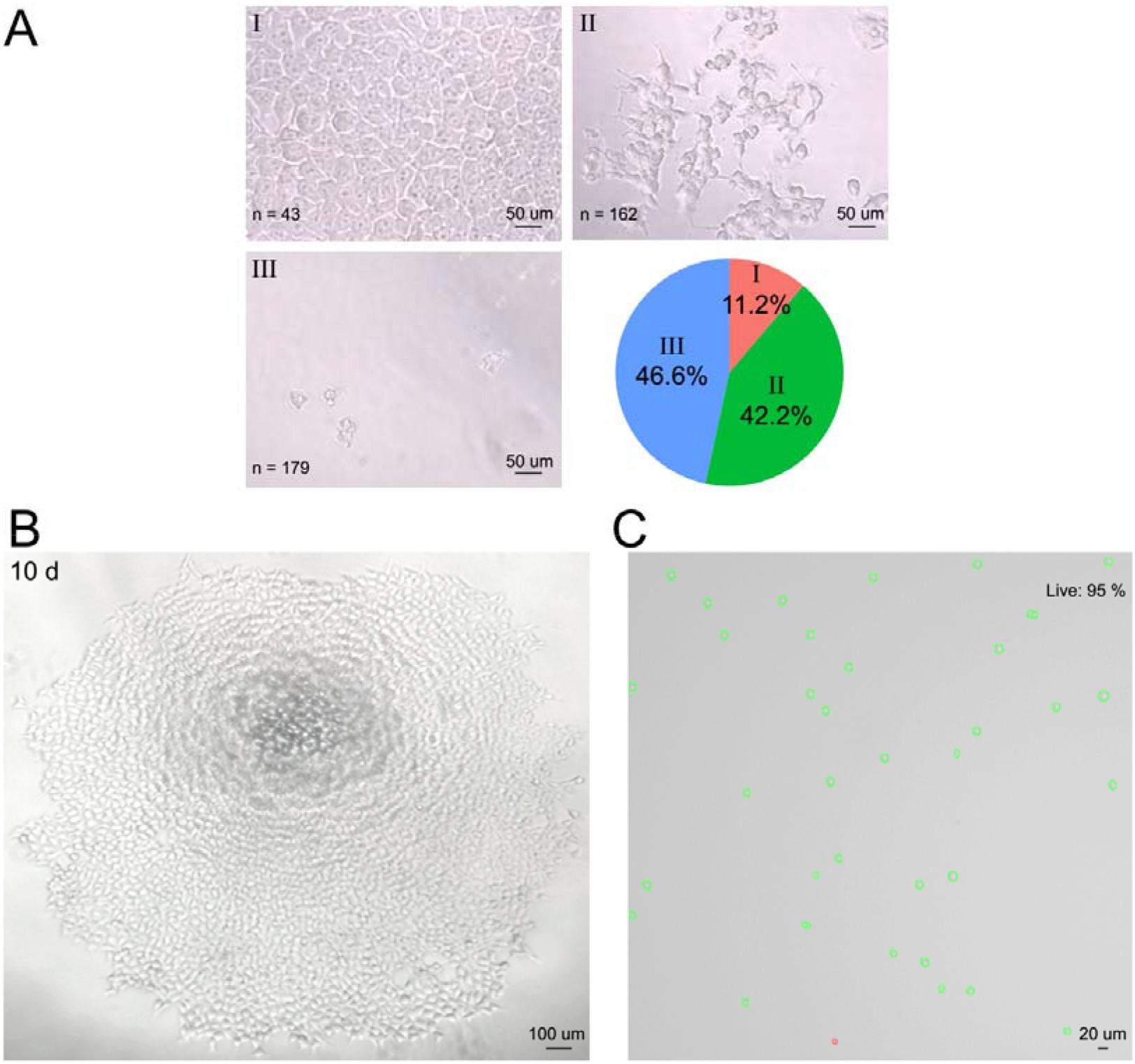
Colony morphology and digestion. (**A**) Typical morphologies of three types (I, II or III) of single-cell colonies. The proportions of the three types of morphologies on the 10^th^ day after seeding with a single cell are shown in a pie chart. The total number of colonies examined was 384. (**B**) The overall morphology of a full type I colony. (**C**) Cell activity of single cells digested from a single-cell colony.

**Figure S4.**
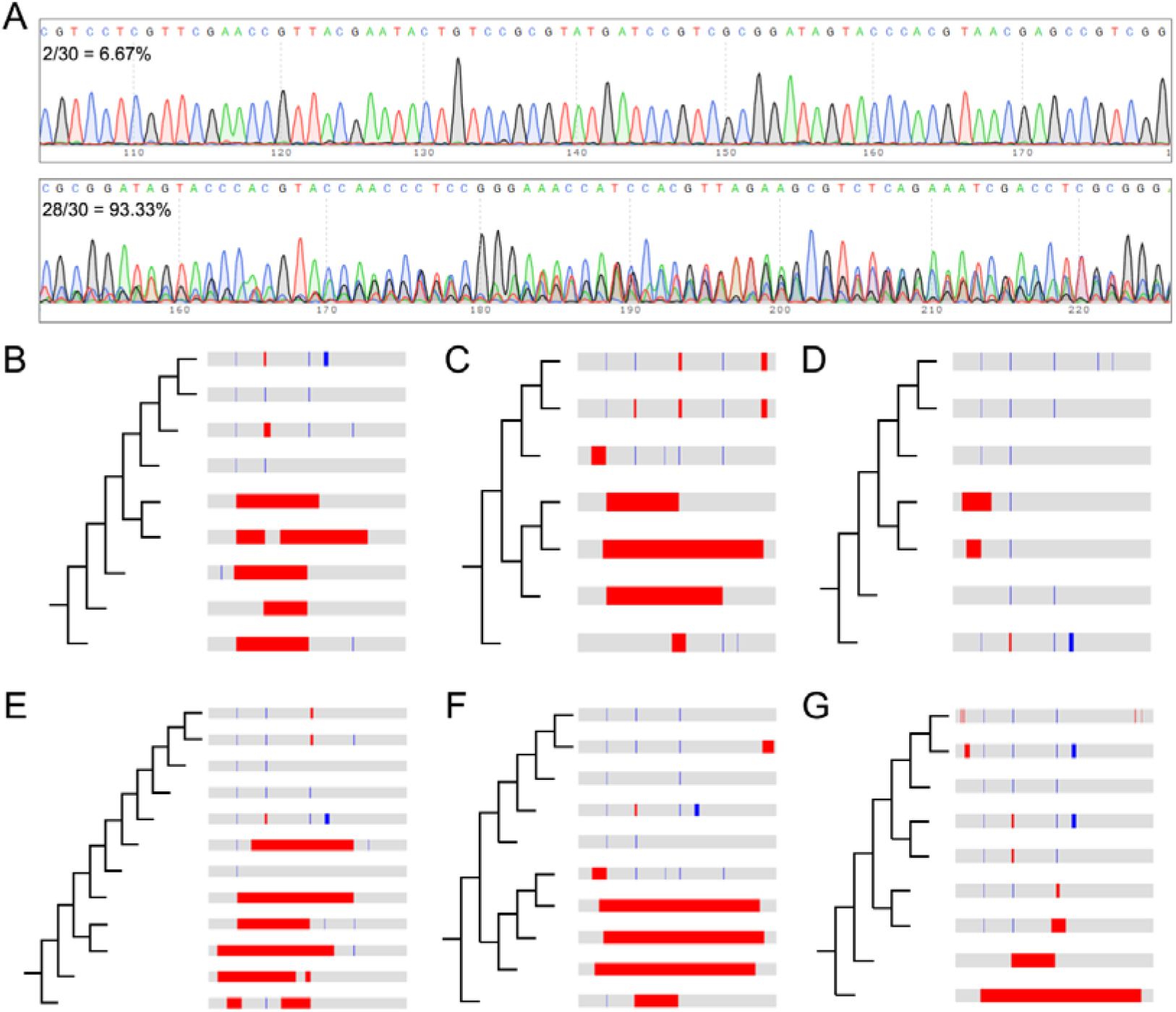
Editing efficiency of the STADT barcodes assessed by TA-cloning followed by Sanger sequencing. (**A**) The editing efficiency of the STADT barcodes in all single-cell clones assessed by the Sanger sequencing of the PCR products was nearly 93%. Two sequencing chromatograms are shown as examples. (**B**-**G**) The diversity of the STADT barcodes in each single-cell clone (transfected with sgRNA lentivirus at different MOIs) cultured with different concentrations of cumate was identified by TA cloning followed by Sanger sequencing (20 clones for each sample). (**B**) sgRNA lentivirus (MOI =10), cumate = 5 µg/ml. (**C**) sgRNA lentivirus (MOI =10), cumate = 30 µg/ml. (**D**) sgRNA lentivirus (MOI =20), cumate = 5 µg/ml. (**E**) sgRNA lentivirus (MOI =20), cumate = 30 µg/ml. (**F**) sgRNA lentivirus (MOI =30), cumate = 5 µg/ml. (**G**) sgRNA lentivirus (MOI =30), cumate = 30 µg/ml.

**Figure S5.**
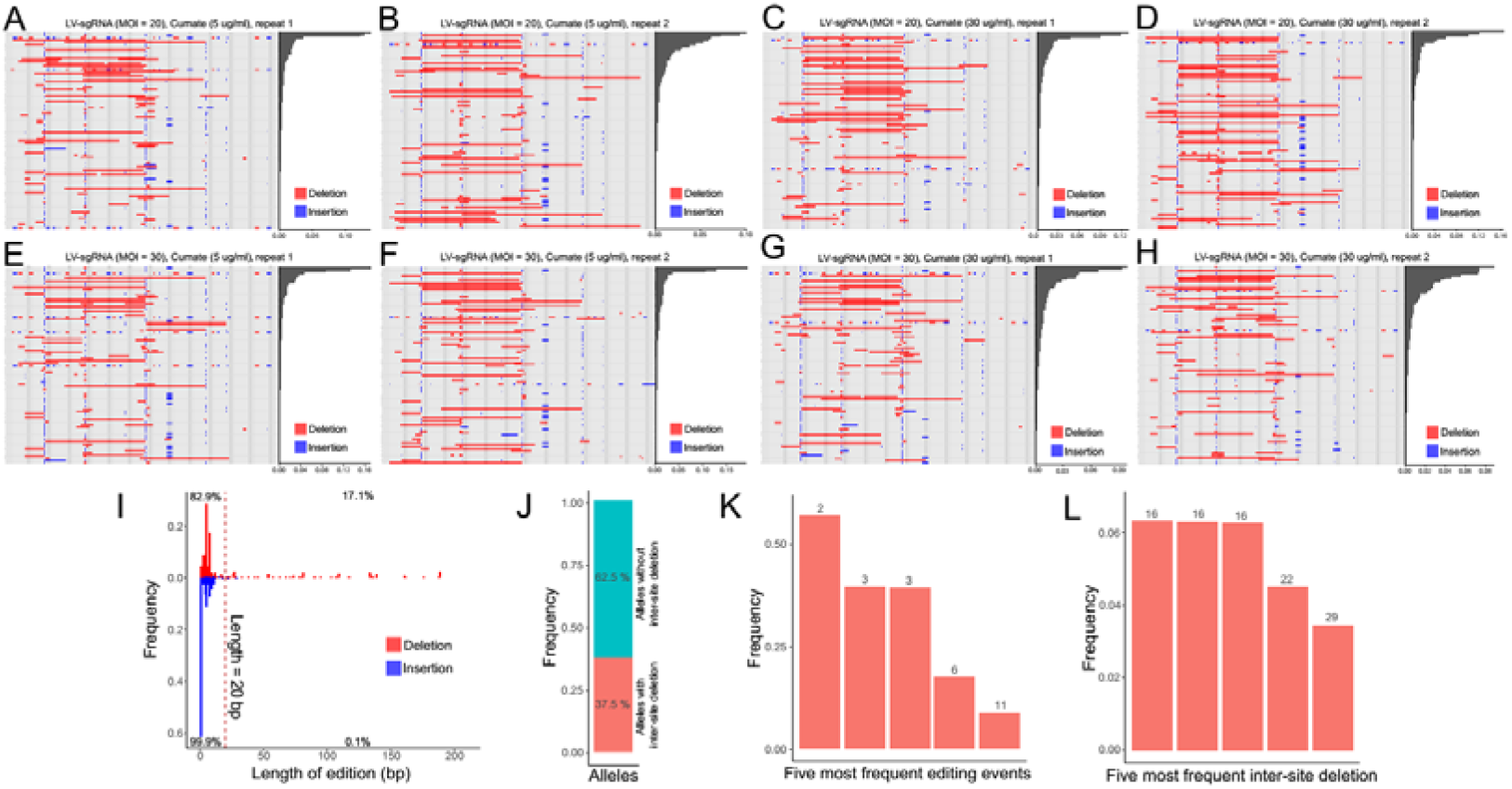
Editing efficiency of the STADT barcodes assessed by HiSeq PE 250 sequencing. (**A**-**H**) The diversity of the STADT barcodes in each single-cell clone assessed by HiSeq-PE250 sequencing. Each panel lists the top 100 most prevalent STADT alleles and their frequencies (the bar plot to the right) detected in a typical colony on the 10^th^ day of growth after seeding with a single cell. The structure of the barcode is color-coded by gray gradients similar to that in Figure 1A. The editing events were indicated by red or blue bars for deletion and insertion, respectively. The MOI of sgRNA lentivirus transfection and cumate concentration during colony growth is indicated on top of each panel. Only the one sample in (**H**) was used in the further analyses below. (**I**) Distribution of the lengths of all editing events. A total of 82.9% of all the deletion events were shorter than 20 bp. (**J**) The frequency of STADT alleles with inter-site deletions. (**K**) Frequency of the top 5 editing events. (**L**) The frequency of STADT alleles with inter-site deletions. For both (**K**) and (**L**), the approximate growth stage (in terms of number of cells) of the single-cell colony in which editing occurred is indicated on top of each bar.

**Figure S6.**
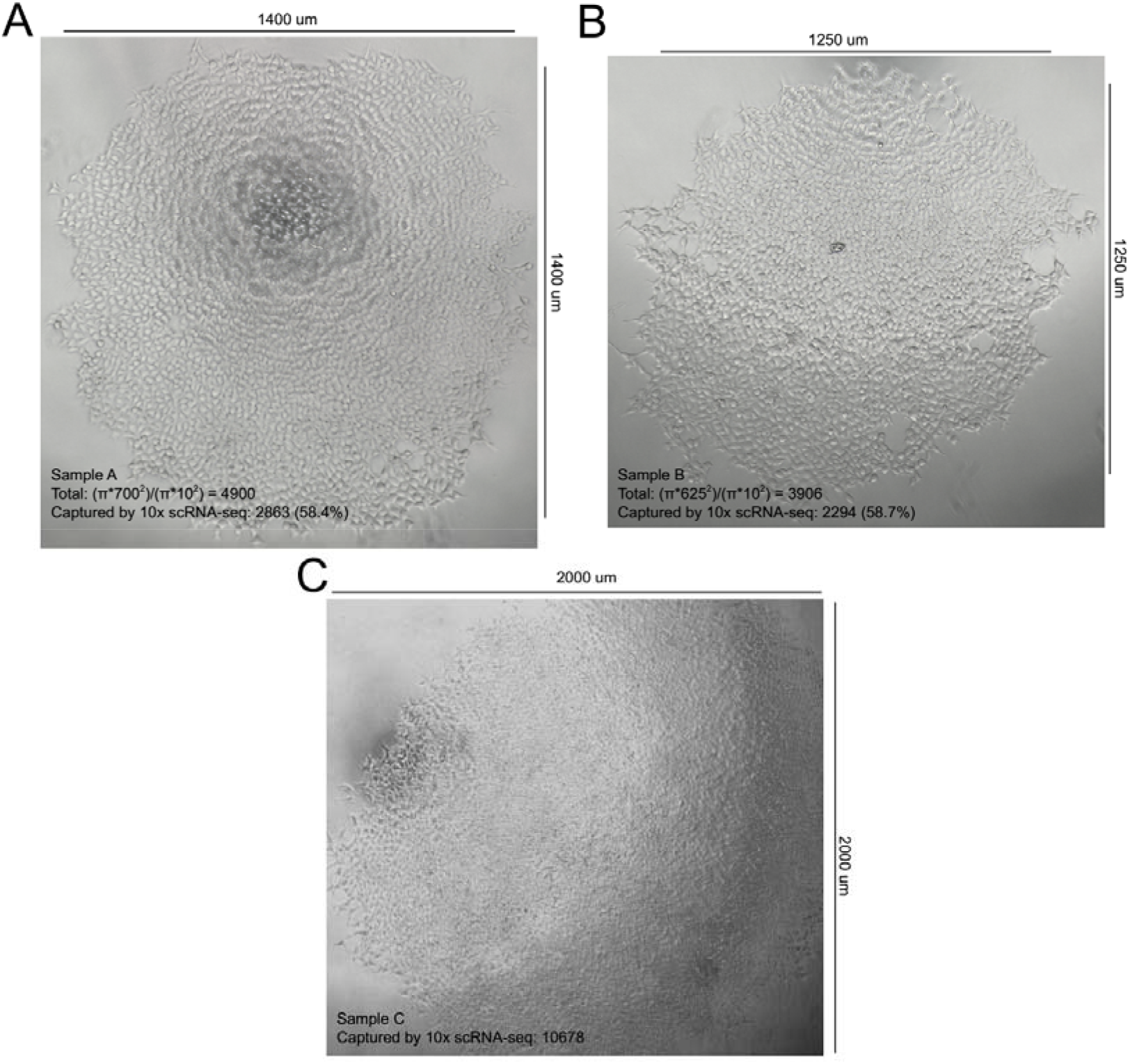
Overall morphologies of single-cell colonies. (**A**-**B**) In colonies with a typical density of cells, the total number of cells was estimated according to the ratio between the area of the colony and the area of a single cell. The values used in approximation are shown at the bottom left corner of each panel, along with the number of cells captured by 10x single-cell RNA-seq. (**C**) A sample with a cell density that was too high to apply the approximation by area-ratio method.

**Figure S7.**
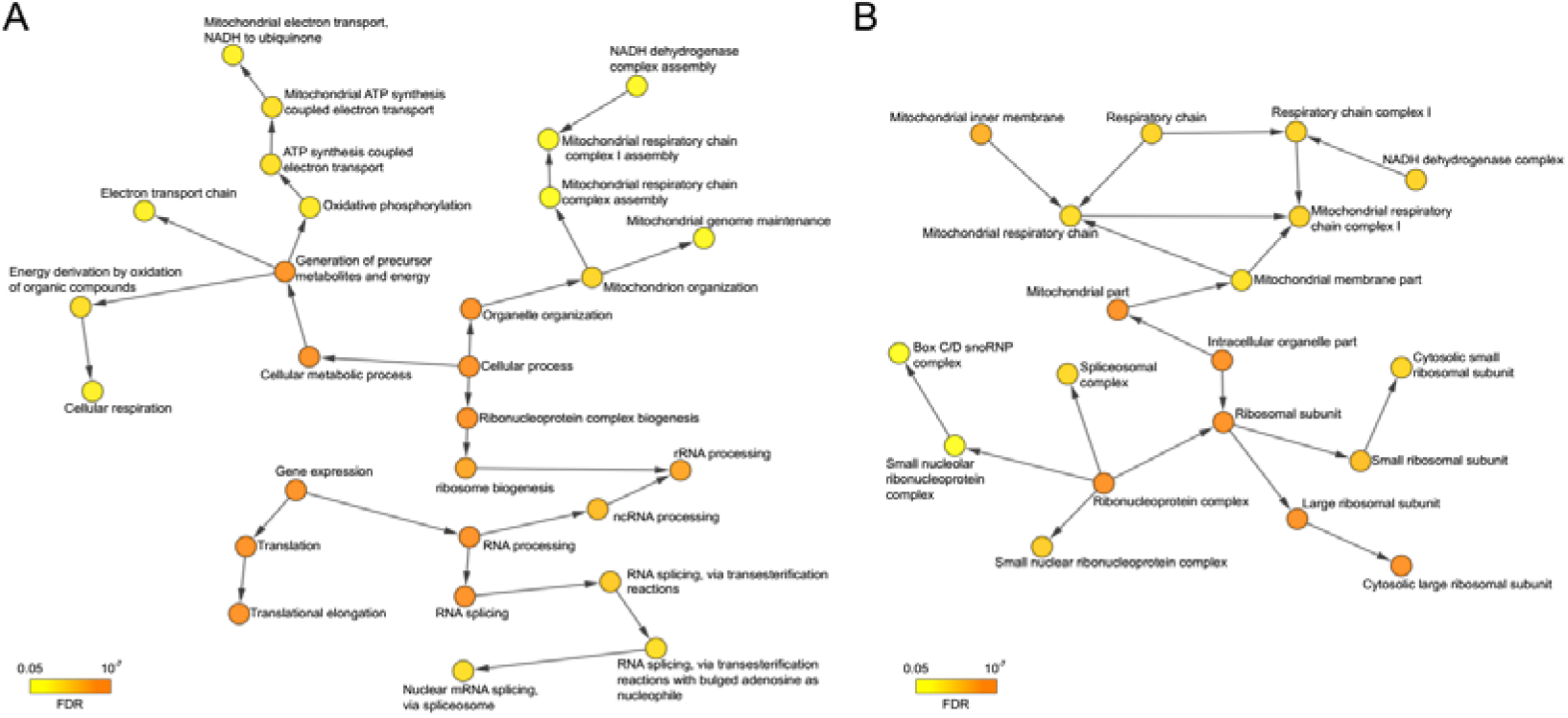
Gene Ontology terms enriched within genes with signature of noise saturation. Functional annotation and GO term enrichment for Biological Process (**A**) and Cellular Component (**B**) for the 492 genes with apparent signature of noise saturation by the BiNGO ^62^ plugin of Cytoscape ^63^ with the default parameters revealed enrichment of mitochondria- and translation/ribosome-related functions.

**Figure S8.**
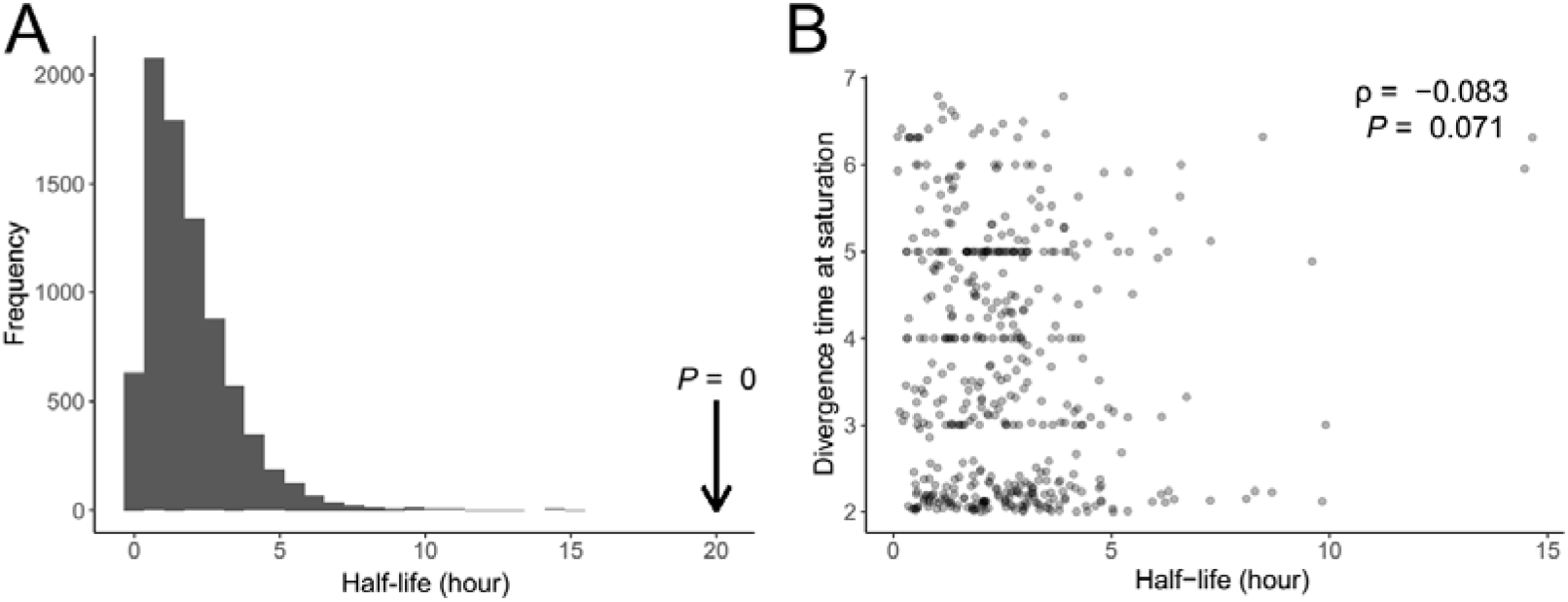
Time of noise saturation cannot be explained by mRNA turnover rate. (**A**) Histogram showing the distribution of mRNA half-lives in HEK-293 cells, as reported previously^31^. The arrow represents the common generation time (duration of a full cell cycle) of HEK-293 cells. The average mRNA half-life and HEK-293 generation time is clearly disparate. (**B**) For each gene, the gene-specific mRNA half-life was compared with the saturation time inferred by the saturation model. The marginally significant negative Spearman’s rank correlation, as indicated on the top-right corner, suggests that the inter-dependence of expression levels among closely related cells cannot be explained by mRNA half-lives.

**Figure S9.**
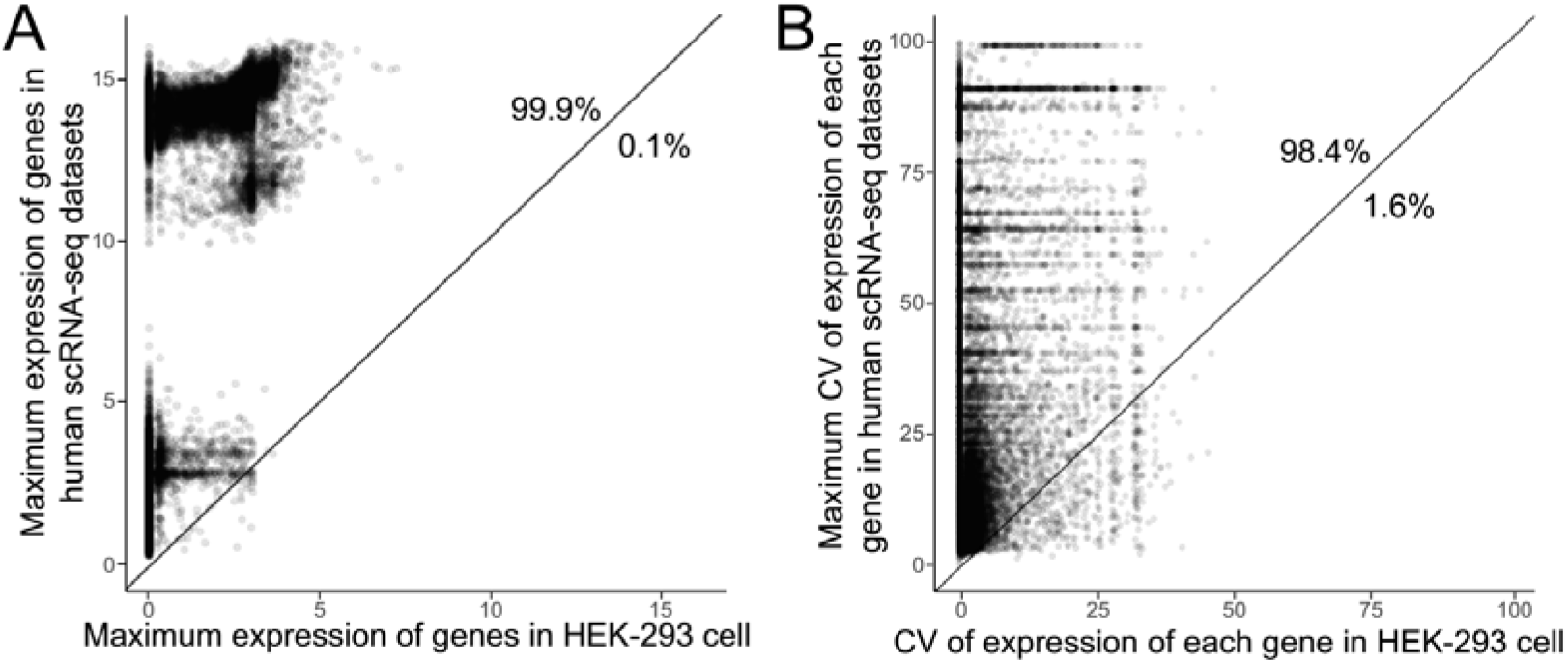
The inferred biological expression boundaries are not physical limits of the expression. (**A**) For each gene, the maximum expression level detected in multiple scRNA-seq datasets^64, 65^ (*x* axis) was compared with that detected in the single cells in our HEK-293 colonies (*y* axis). Most (99.9%) genes are above the diagonal line (*x* = *y*), suggesting that the physical maximum of their expression level was not reached. (**B**) For each gene, the maximum CV of the expression level estimated in multiple scRNA-seq datasets ^64, 65^ (*x* axis) was compared with the CV estimated in our HEK-293 colony (*y* axis). Most (98.4%) genes are above the diagonal line (*x* = *y*), suggesting that the physical maximum of their expression noise was not reached.

