## Supplementary figures and images for "A high-density lineage tree reveals dynamics of expression differences accumulation in nondifferentiating clonal expansion"

### Supplemental Figures

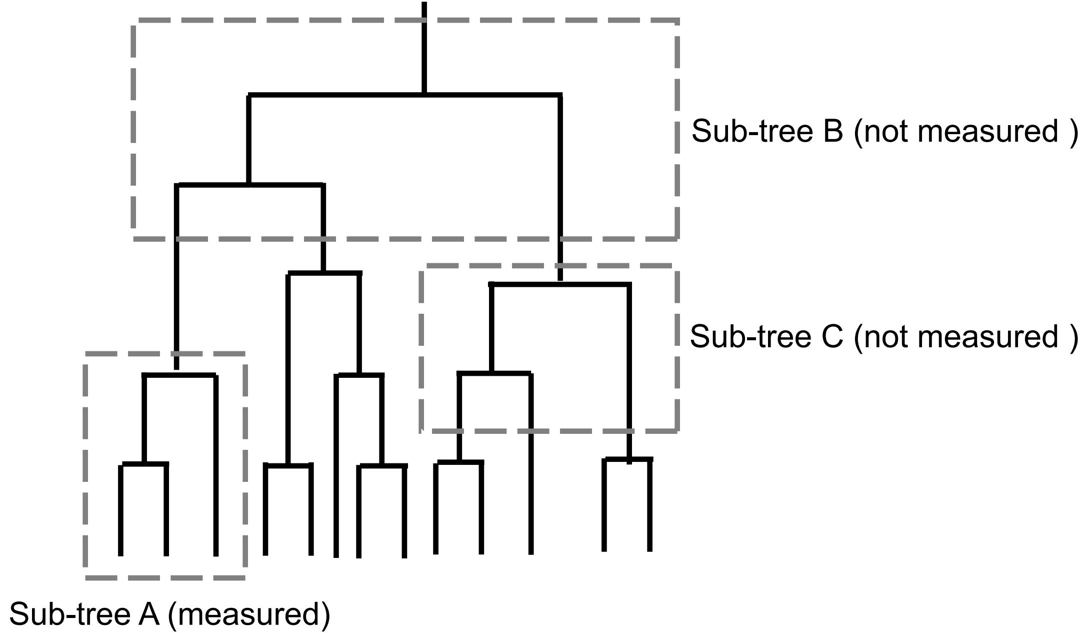

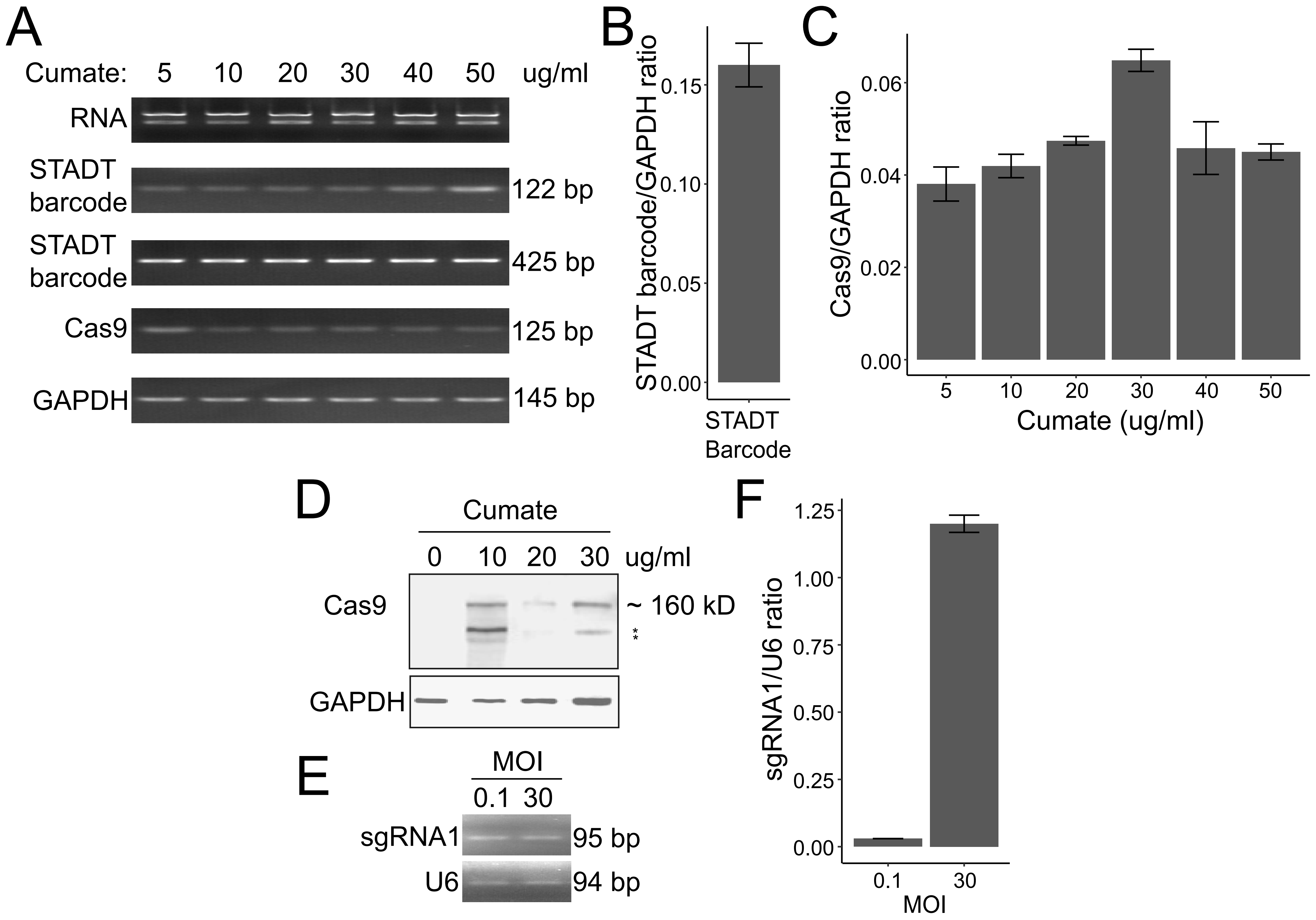

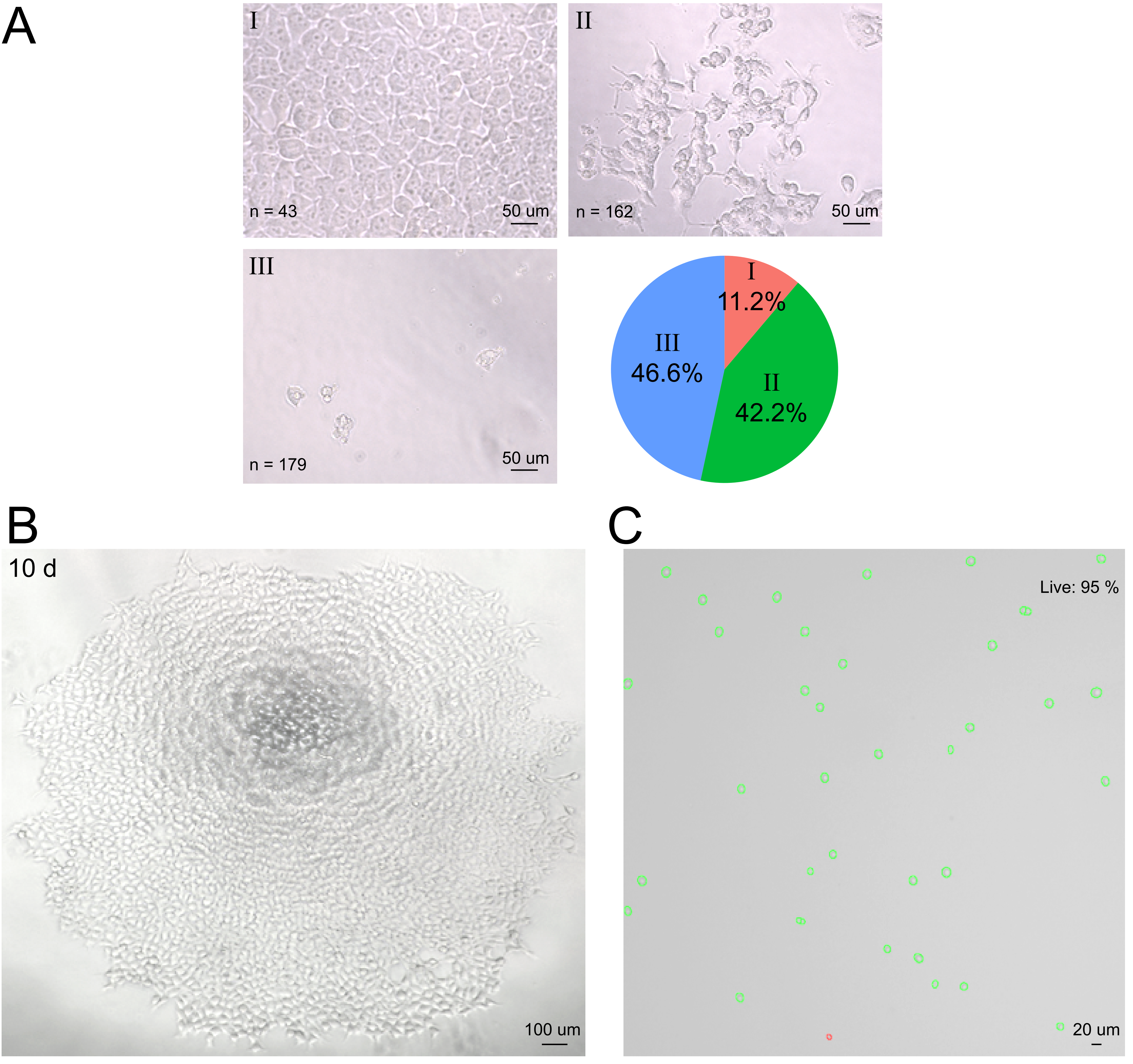

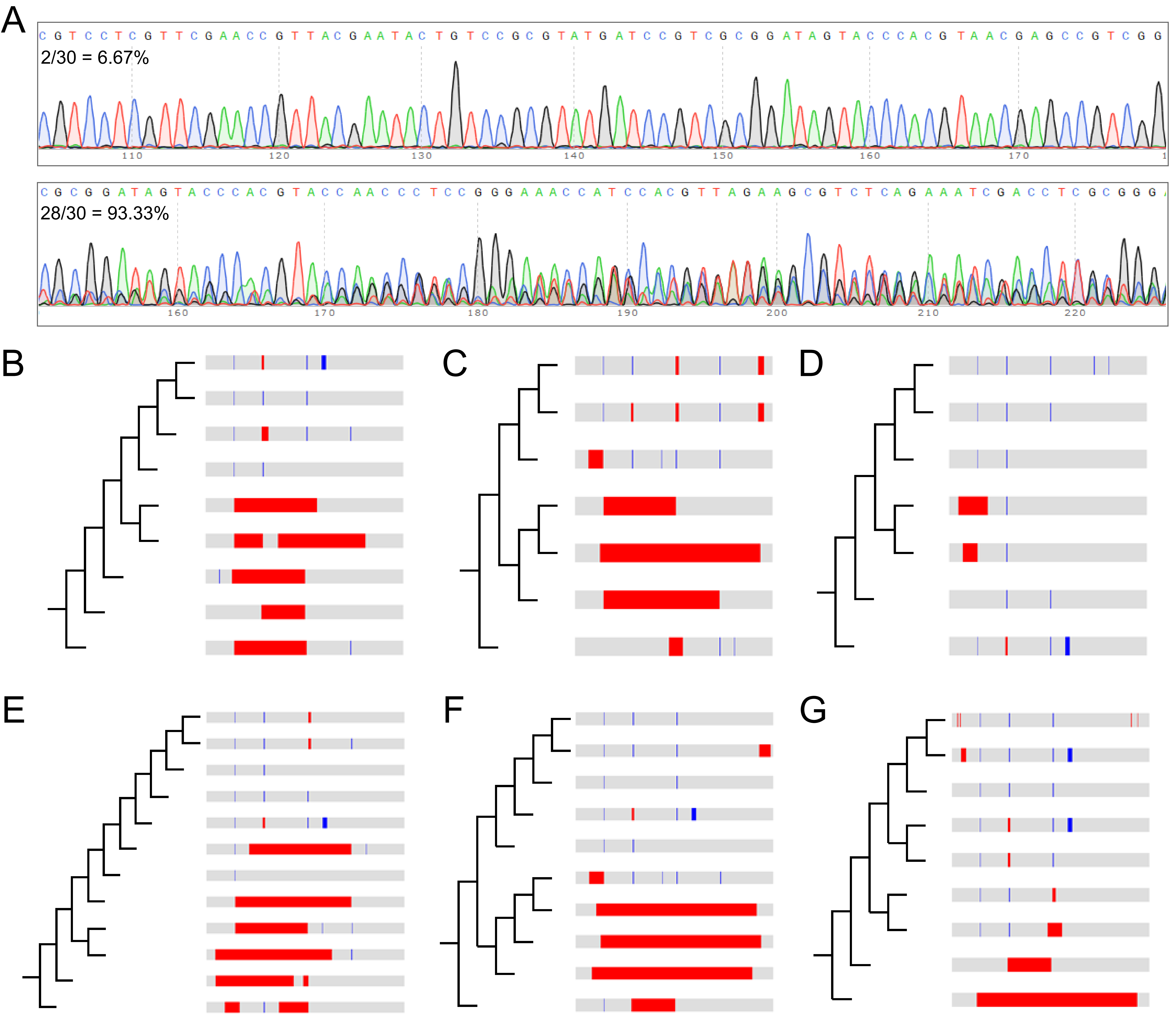

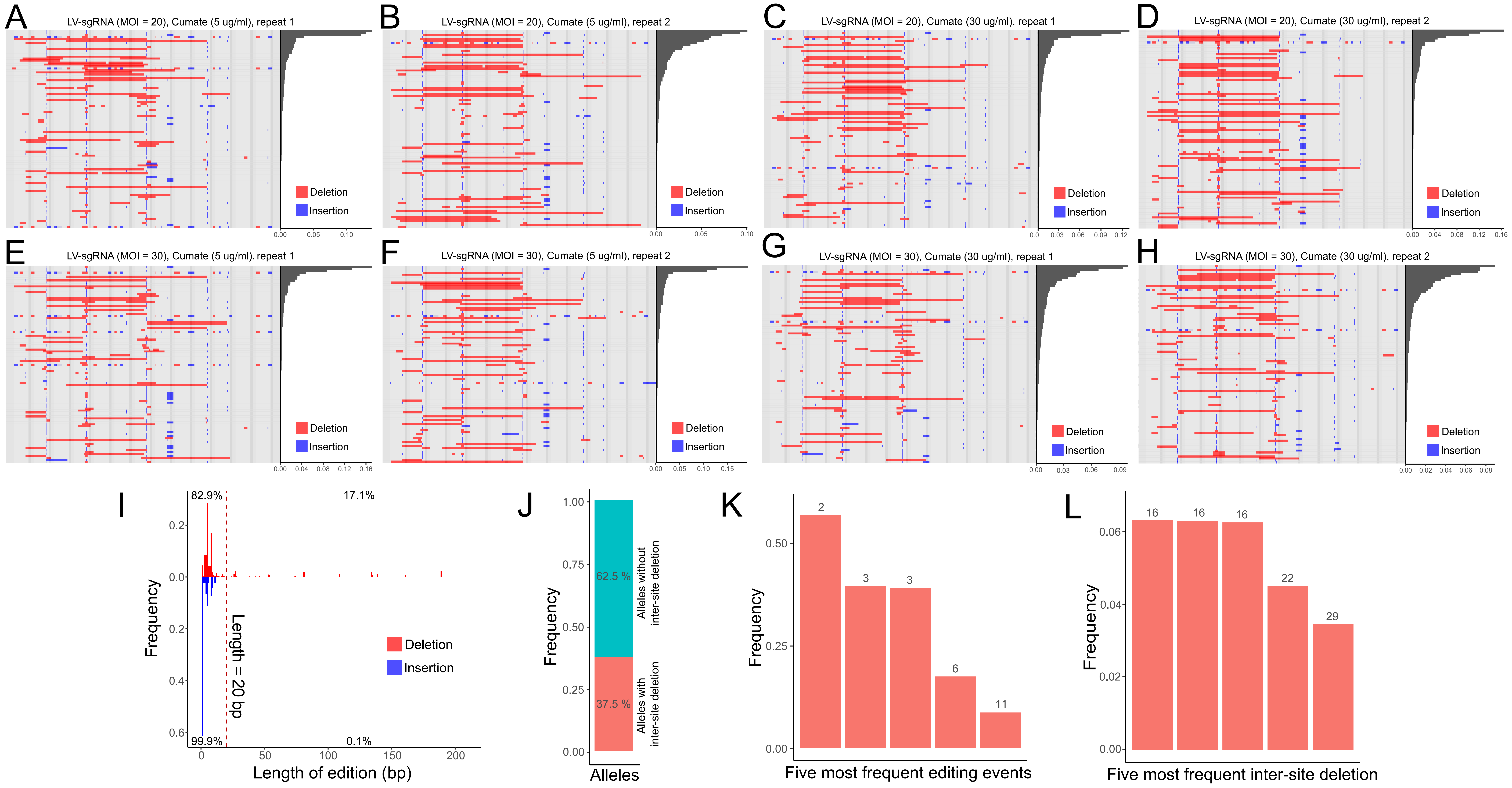

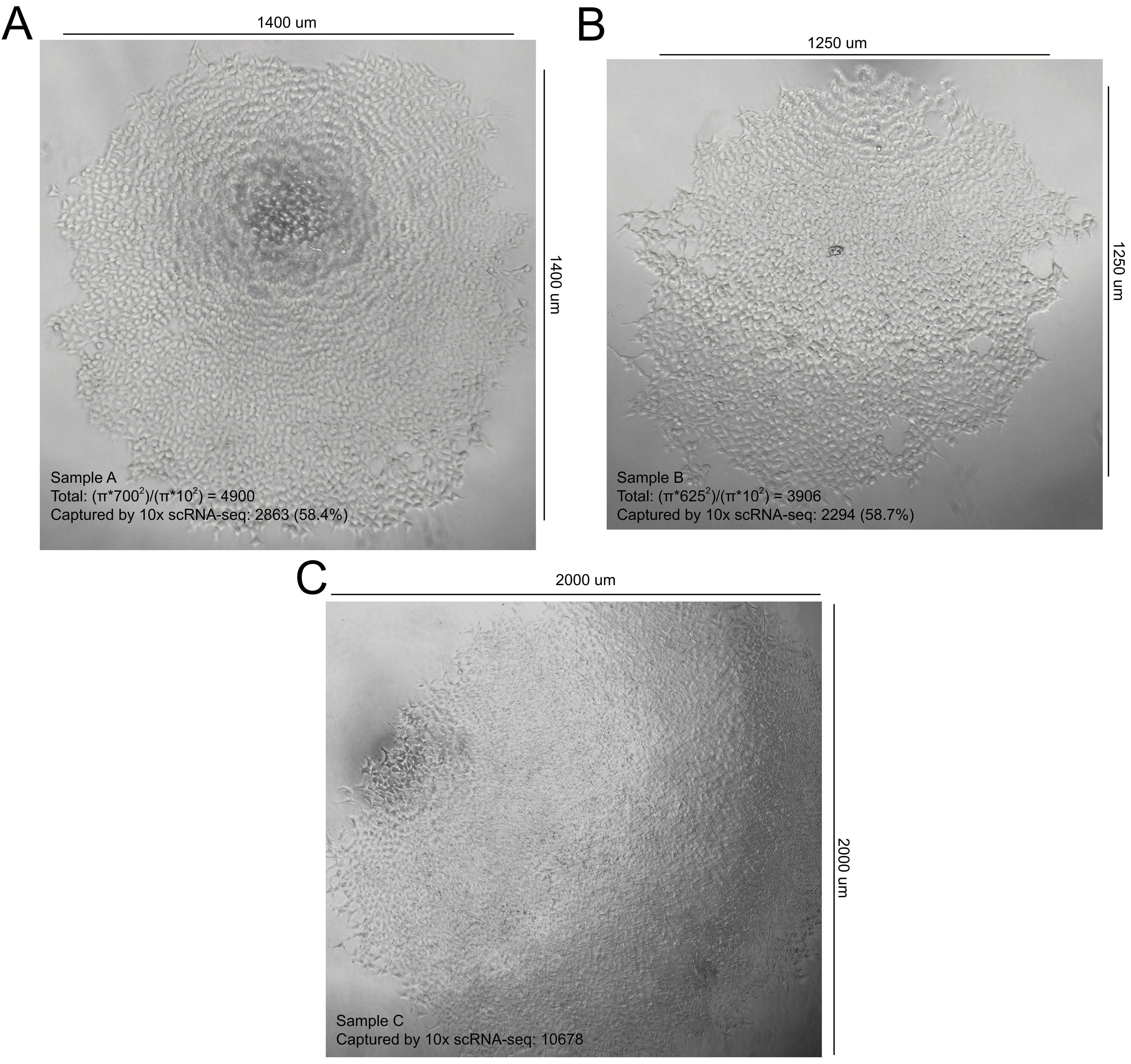

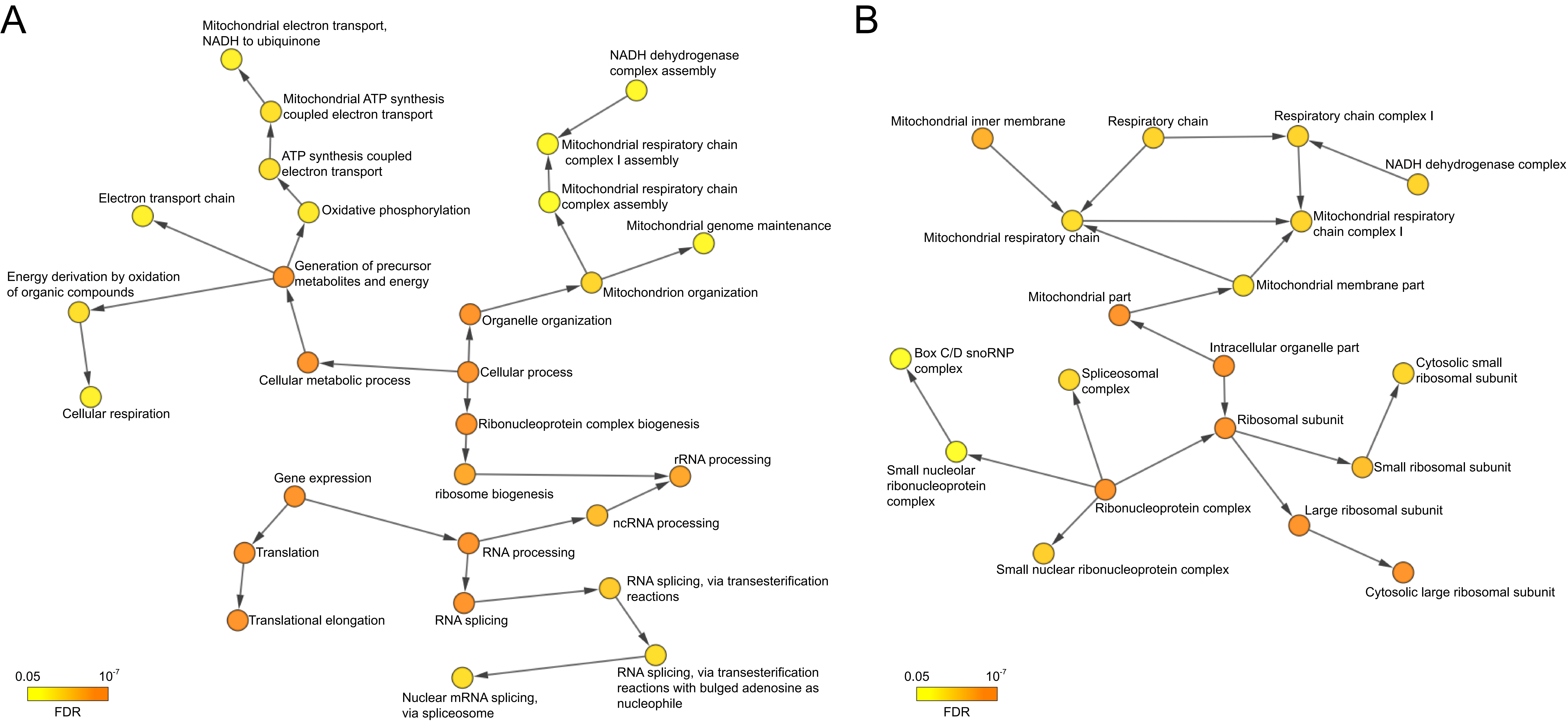

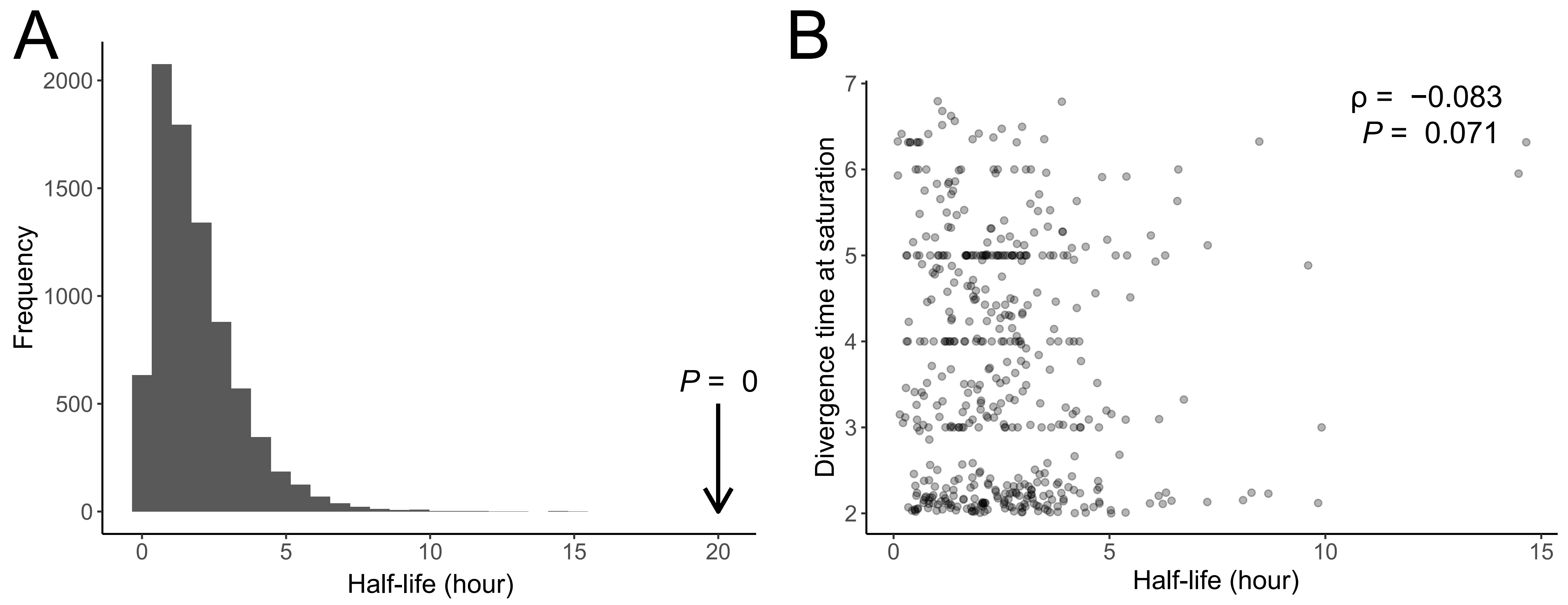

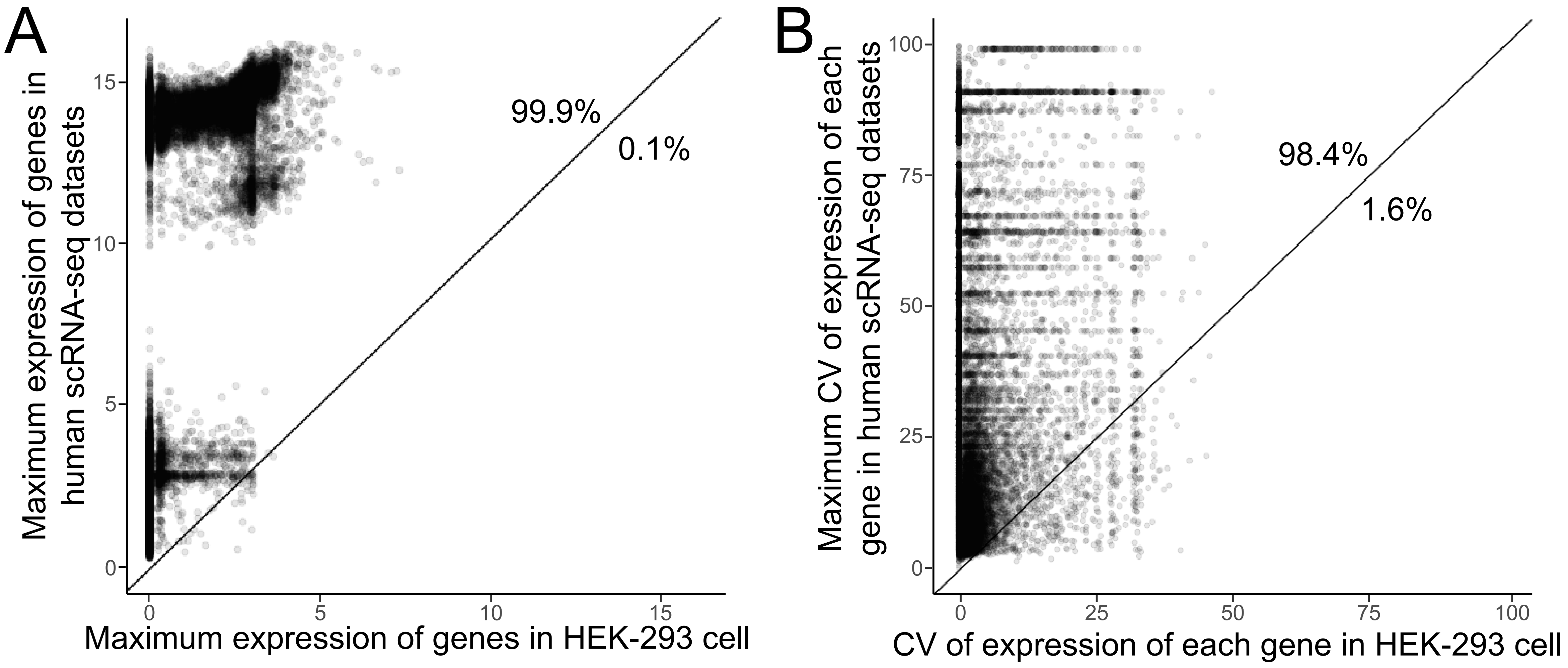
